# Differences in nanoscale organization of DNase I hypersensitive and insensitive chromatin in single human cells

**DOI:** 10.1101/2021.05.27.445943

**Authors:** Katharina Brandstetter, Tilo Zülske, Tobias Ragoczy, David Hörl, Eric Haugen, Eric Rynes, John A. Stamatoyannopoulos, Heinrich Leonhardt, Gero Wedemann, Hartmann Harz

## Abstract

Methodological advances in conformation capture techniques have fundamentally changed our understanding of chromatin architecture. However, the nanoscale organization of chromatin and its cell-to-cell variance are less studied. By using a combination of high throughput super-resolution microscopy and coarse-grained modelling we investigated properties of active and inactive chromatin in interphase nuclei. Using DNase I hypersensitivity as a criterion, we have selected prototypic active and inactive regions from ENCODE data that are representative for K-562 and more than 150 other cell types. By using oligoFISH and automated STED microscopy we systematically measured physical distances of the endpoints of 5kb DNA segments in these regions. These measurements result in high-resolution distance distributions which are right-tailed and range from very compact to almost elongated configurations of more than 200 nm length for both the active and inactive regions. Coarse-grained modeling of the respective DNA segments suggests that in regions with high DNase I hypersensitivity cell-to-cell differences in nucleosome occupancy determine the histogram shape. Simulations of the inactive region cannot sufficiently describe the compaction measured by microscopy, although internucleosomal interactions were elevated and the linker histone H1 was included in the model. These findings hint at further organizational mechanisms while the microscopy-based distance distribution indicates high cell-to-cell differences also in inactive chromatin regions. The analysis of the distance distributions suggests that direct enhancer-promoter contacts, which most models of enhancer action assume, happen for proximal regulatory elements in a probabilistic manner due to chromatin flexibility.

## Background

For almost one hundred years it has been known that interphase chromatin can be distinguished by means of light microscopy into less dense euchromatin and denser packed heterochromatin (1, 2). Later it became clear that nucleosomes are the basic building blocks organizing DNA packaging and are therefore central to the organization of chromatin (3). Groundbreaking electron microscopic studies showed the tight interaction between nucleosomes and DNA forming an 11 nm thick fiber (4, 5). Recent work reveals a more random, heterogeneous organization of chromatin (6, 7). This view is supported by electron microscopic studies and super-resolution fluorescence microscopy that show interphase chromatin to be organized in a flexible and disordered structure where regions with higher nucleosome density are interspersed with nucleosome depleted regions (8–11).

The landscape of chromatin states is much more diverse than the originally described eu- and heterochromatin suggest. By analyzing genome-wide distribution patterns of chromatin associated proteins, posttranslational histone modifications and DNase I hypersensitivity with algorithms like ChromHMM and Segway, up to 51 chromatin classes were proposed (12–18). DNase I hypersensitivity (DHS) is a criterion that can also be used alone to subdivide chromatin in regulatory or active DNA with high DHS as opposed to inactive regions with low DHS (19, 20).

Posttranslational histone modifications of the active chromatin classes, like acetylation, usually reduce nucleosome interaction strength and thus produce an open, less densely packed chromatin (21–25). Inactive classes are often characterized by methylation marks on histone 3 (e.g. H3K9me2/3), which can be bound by the heterochromatic protein 1 (HP1), thereby compacting chromatin (26). However, large parts of inactive and more densely packed chromatin do not carry significant amounts of posttranslational histone modifications (12). Other mechanisms must therefore be responsible for compaction.

A remarkable feature of chromatin is its dynamic and fluid nature which has been observed in several fluorescence imaging studies (27–37) and is the reason for the large cell-to-cell variability in the structure of chromatin domains (38). Changes in nucleosome occupancy are actively regulated and can drastically affect the 3D genome architecture as it has been shown e.g. by the effects of tumor necrosis factor alpha on human endothelial cells (39). Even at the level of single nucleosomes, a significant and dynamic cell-to-cell variability can be found (40). The recently developed Fiber-seq method reveals that regulatory elements are actuated in an all-or-none fashion, thereby replacing a canonical nucleosome (41). In addition to pioneer transcription factors, some chromatin remodelers are known to exhibit nucleosome eviction activity (42–44). Together, these examples show that, depending on the regulatory context, the number and exact position of nucleosomes in active chromatin of eukaryotes can dynamically change.

Computational studies show a close link between nucleosome positions and the spatial organization of chromatin (45) which was explored by applying coarse-grained computer simulations by many groups (e.g. (46–48)). These studies demonstrate, for example, that different nucleosome repeat lengths are responsible for more open or closed chromatin configurations (49). Moving even a single nucleosome can strongly influence the spatial organization (50). Thus, including the real length of the different linker DNA into coarse-grained models is required to obtain realistic results (50).

In our research, we investigated structural differences between oligoFISH-labeled active and inactive 5 kb chromatin segments of prototypical chromatin regions, selected on the basis of the presence or absence of DNase I hypersensitivity. By measuring the distance between labeled endpoints with systematic 3D STED microscopy and comparing this data with coarse-grained Monte Carlo simulations (50, 51) we aimed to find underlying organizational principles. In active chromatin, simulated data match the microscopic data well, assuming cell-to-cell variability in nucleosomal density. For inactive chromatin, the fit between model and microscopic measurements was generally lower, indicating additional compaction mechanisms that act in parallel to increased internucleosomal energy and the presence of the linker histone H1. Regardless of whether chromatin is active or inactive, our results reveal two striking features for 5 kb segments: (i) all distance distributions are right-tailed, and simulations indicate an underlying cell-to-cell variance in chromatin organization, (ii) distributions cover a wide range of distances from less than 50 nm to more than 200 nm.

## Results

Chromatin organization of active and inactive chromatin was analyzed in K-562 cells using systematic super-resolution microscopy of DNA sequences labeled with oligoFISH probes and comparison with simulated 3D chromatin configurations generated by a coarse-grained model. The K-562 cell line is well suited for computer simulations as a wealth of information like genome-wide ChIP-seq data, comprehensive maps of posttranslational nucleosome modifications and nucleosome positioning generated by the ENCODE project are available (15, 52).

### STED microscopy as a tool to study prototypic chromatin regions on the kb scale

By using data from the ENCODE project we selected a 20 kb region on chromosome 11 (hg19, chr11: 118955404 - 118977871) which exhibits very high hypersensitivity to DNase I not only in K-562 (Fig. 1 a), but also in more than 150 other cell types. Moreover, this region is flanked up- and downstream by highly active chromatin. For inactive chromatin, the selection criteria were analogous: missing DNase I hypersensitivity over 30 kb in 651 investigated cell types with over 2 Mb without DHS in K-562 cells. The selected 20 kb inactive region is also located on chromosome 11 (hg19, chr11: 55580425 - 55603312) (Fig. 1 b). For each of these 20 kb regions 5 oligoFISH probe sets (A, B, C, D, E; Fig. 1 a,b) were designed, dividing the 20 kb into four approximately 5 kb long segments from midpoint to midpoint of the respective probe set (probe set combinations: AB, BC, CD, DE). Each oligoFISH probe set consisted of 30 oligonucleotides (directly fluorescently labeled 40mers) covering a region of about 1.5 - 2 kb (Fig. 1 a, b). These small genomic distances are expected to result in spatial distances falling below the resolution limit of light microscopy (53) which is about 250 nm in the x- and y-dimensions and more than 500 nm in z (54). Two-color super-resolution 2D and 3D STED microscopy was employed to overcome this limitation. STED microscopy is not prone to any chromatic shift if (present case) the different fluorophores are depleted by the same doughnut (55). The two-color approach also allows the use of subpixel localization techniques to measure distances below the resolution limit of the STED microscope.

**Fig. 1:**
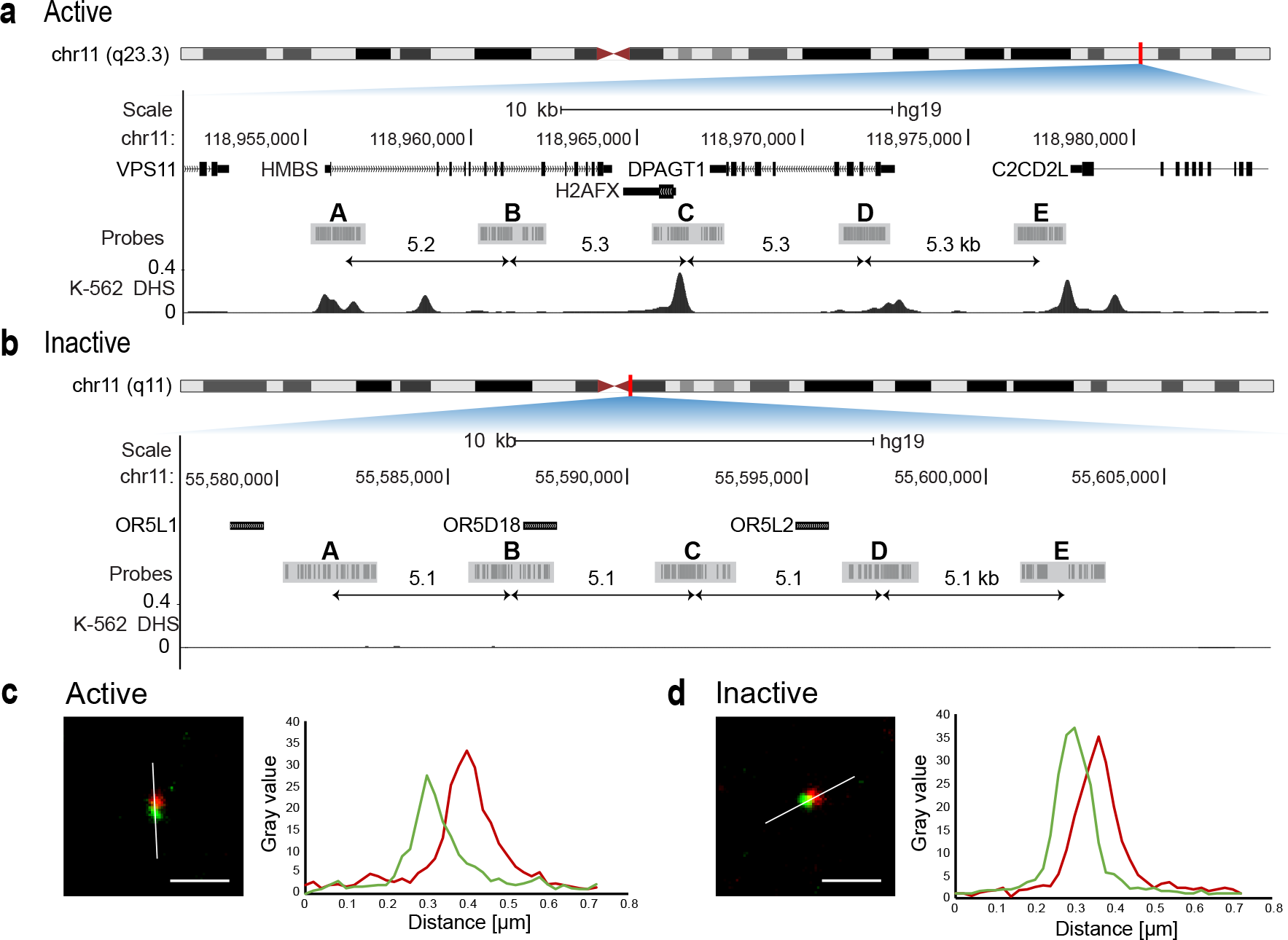
FISH probe design for active and inactive region. Both regions are located on chromosome 11. (a) The active region contains genes *HMBS*, *H2AFX* and *DPAGT1*. The probe sets are almost equally spaced (5.2, 5.3, 5.3, 5.3 kb midpoint to midpoint) and mostly cover DNase I hypersensitive sites. (b) The inactive region contains genes for olfactory receptors. The region shows no DNase I hypersensitivity and the probe sets are equally spaced (5.1 kb midpoint to midpoint). (c, d) Example images show STED detail images of FISH spots in two colors for active (c) and inactive (d) (Target 1 in green, target 2 in red). Measured distance of these shown spot pairs represents the mean of the population. Plots depict intensity values for both colors along lines of interest (white lines). Scale bar = 500 nm

### Inactive regions are more compact than active regions

Recent studies reveal a high cell-to-cell variance of the spatial genome organization (56–58). To study the chosen regions, we applied high-throughput 2D STED microscopy to generate data with high statistical power characterizing the nanoscale organization of 5 kb segments of active and inactive chromatin. For each of the 8 investigated 5 kb segments between 484 and 1621 single cell measurements were analyzed. The four measured intervals in the active chromatin region differ from one another. We found some significant deviations with the maximum difference in the median projected distance of 16 nm (p=0.00053, BC versus DE and CD versus DE, Wilcoxon rank sum test) (Fig. 2a, Supplementary Table 1). In active chromatin, variability of the nanoscale organization is expected since each 5 kb segment is composed of different proportions of exons, introns, enhancers and other regulatory sequences. Surprisingly, we also found highly significant differences between the investigated intervals in inactive chromatin. We expected much less difference in compaction because inactive chromatin is expected to be more uniform as it does not harbor active regulatory elements and nucleosome occupancy is not modified by transcriptional activity (Fig. 2b, Supplementary Fig. 1). The maximum difference in the median projected distance was 12 nm within the inactive chromatin group (p<0.0001, AB versus DE, Wilcoxon rank sum test, Supplementary Table 1).

**Fig. 2:**
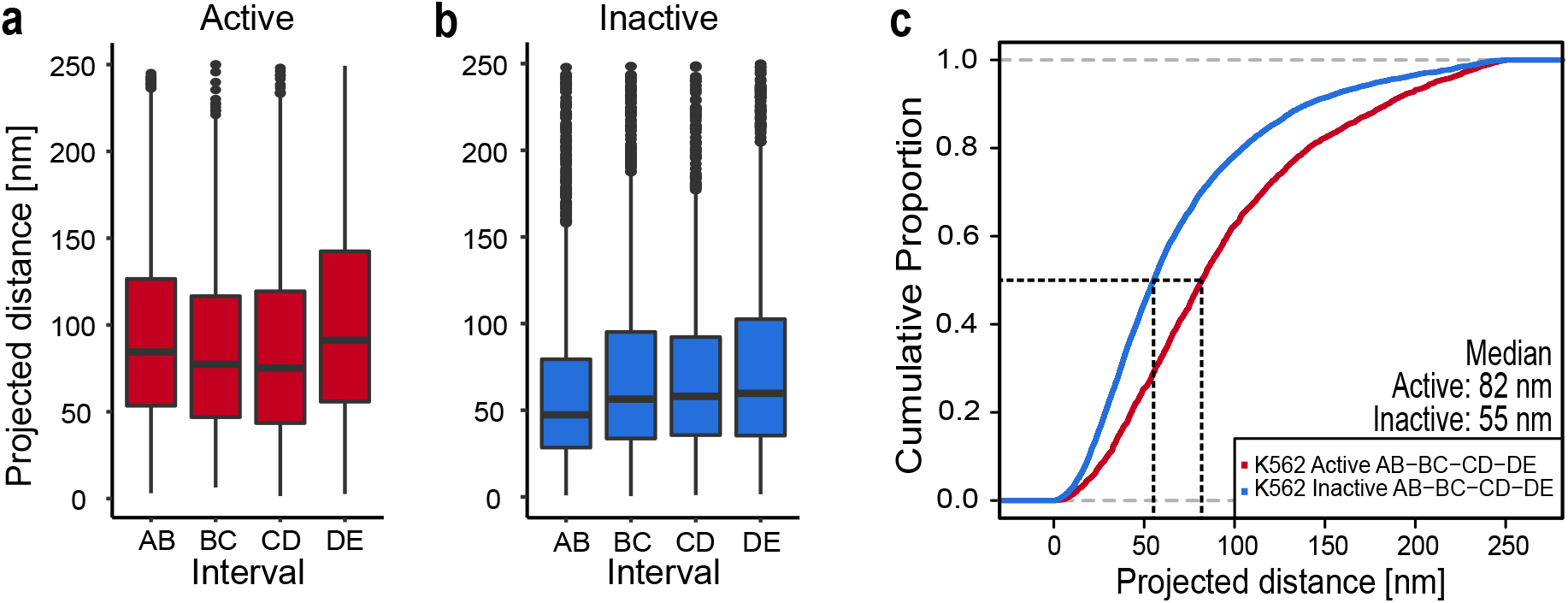
2D STED distance measurements showed that the inactive region is more compact than the active region. (a) Boxplot for the active region for all four measured intervals (AB: n=672, BC: n=540, CD: n=484, DE: n=566, n=number of single-cell measurements pooled from three independent replicates). (b) Boxplot for the inactive region for all four measured intervals (AB: n=1585, BC: n=1621, CD: n=1200, DE: n=1395, n=single-cell measurements from three independent replicates). (c) All data from active (a) and inactive (b) were pooled to generate a cumulative distribution. The cumulative distribution of measured distances showed differences in distributions between active (red) and inactive (blue). The median is the value at the 50 % proportion (black dashed line). For the active region the median is 82 nm, for the inactive region it is 55 nm.

However, since the differences within the active and inactive regions are small, they were pooled to show the overall length distribution of each chromatin class. The median projected distance between two FISH spots flanking a typical 5 kb interval of active chromatin is 82 nm, and 55 nm in inactive chromatin (Fig. 2 c). Shorter double spot distances indicate a higher degree of chromatin compaction whereas larger distances suggest less compaction. Thus, data from our measurements are in line with published data showing active chromatin to be less compacted compared to inactive chromatin (59). As expected, the distributions of the FISH spot distances of active and inactive chromatin differ significantly as shown in a cumulative distribution plot (Fig. 2 c, p<2×10^−16^, Wilcoxon rank sum test, Supplementary Table 1).

For a more in-depth analysis we selected a 5kb segment for both the active and inactive region which are representative of the respective group in 2D STED measurements. We chose interval AB for the active region and CD for the inactive region (Fig. 2 a, b).

### Assigning the input parameters for coarse-grained modeling

The exact position of nucleosomes is an important input parameter for coarse-grained models and strongly affects simulated configurations (50). Nucleosomal positioning can be determined by micrococcal nuclease digestion followed by deep sequencing (MNase-seq) (60). Here we used ENCODE MNase-seq tracks of K-562 cells which are derived from cell populations and therefore often show a seemingly overlapping nucleosome pattern (UCSC Accession: wgEncodeEH000921, GEO Accession: GSM920557). These data are unsuitable for our coarse-grained model, as it requires non-overlapping unique nucleosome positions as input. Therefore, we computed the most probable non-overlapping nucleosome populations by applying the NucPosSimulator (61). Experimentally derived nucleosome occupancy and the computed most probable nucleosome positions of active region AB and inactive region CD are shown in Fig. 3 a and b. Nucleosome positions of the respective flanking regions can be found in Supplementary Fig. 1. For the nucleosomal repeat length (NRL) of chromosome 11 we calculated a mean value of 183.4 +/− 66.3 bp applying NucPosSimulator (Fig. 3 c) (for calculation details see Methods section). The mean NRL of the active (AB) and inactive (CD) region studied in detail is 179.6 bp and 179.1 bp, respectively (Fig. 3 d). Both values are in the range of the NRL of chromosome 11.

**Fig. 3:**
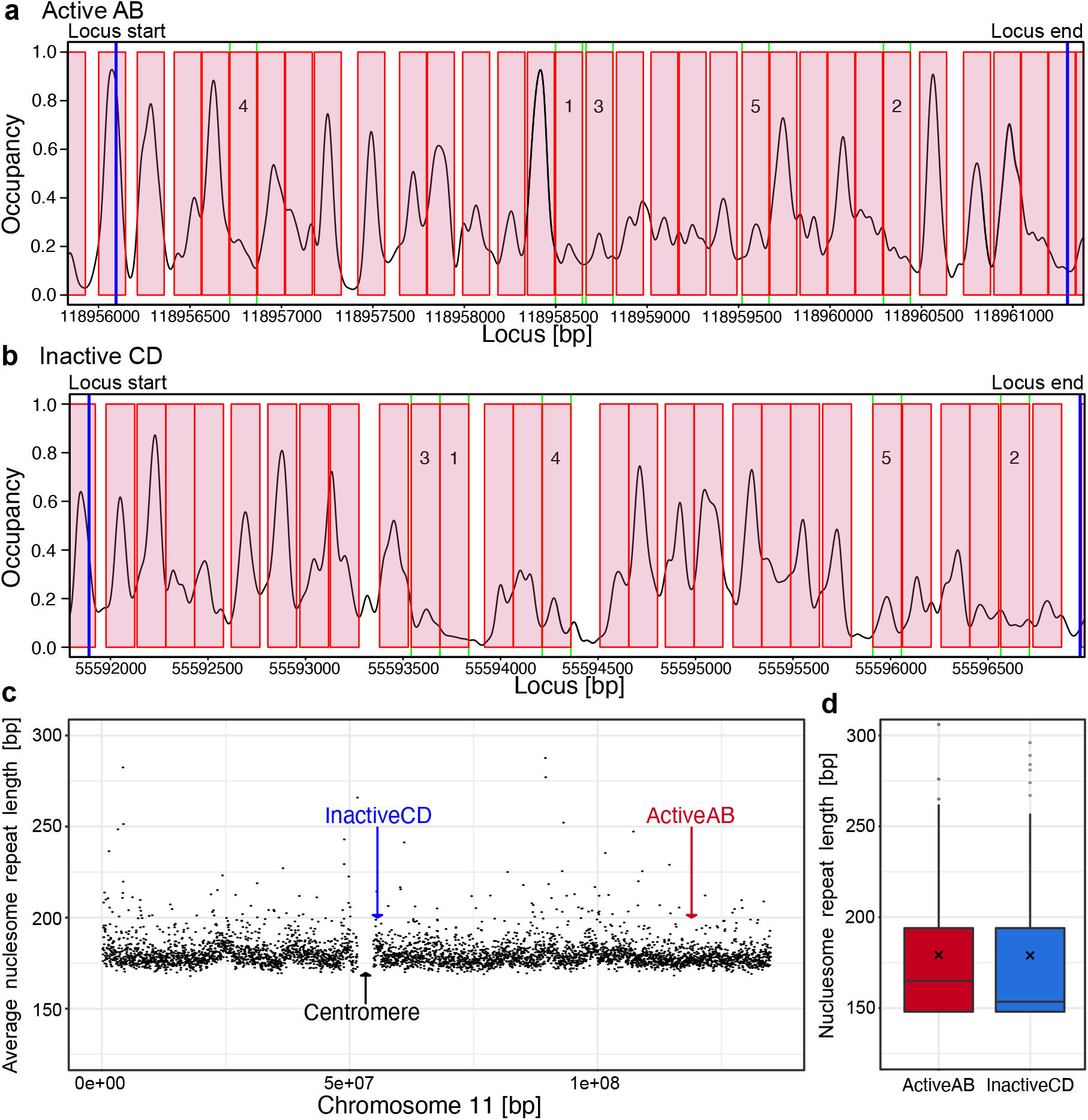
Nucleosome positions and nucleosome repeat length were calculated using the NucPosSimulator. Nucleosome positions (red boxes) for Active AB (a) and Inactive CD (b) based on MNase-seq occupancy tracks (black line). Blue lines indicate start and end of the investigated loci. Numbers in boxes indicate the ranking of the 5 nucleosomes with the lowest binding probability. (c) Mean values of the NRL of a sliding window of the size 30000 bp. Values larger than 300 and windows with fewer than 3 nucleosomes were omitted. The mean NRL for chromosome 11 was 183.4 +/− 66.3 bp. (d) Investigated active and inactive regions as marked in the plot (arrows in c) have a mean of 179.6 bp and 179.1 bp, respectively (black x).

The internucleosomal energy is another important parameter in all coarse-grained models and depends on the solvent (62) and histone modification (58). Literature values for this energy range from 3 to 10 kT (58, 63, 64). Nucleosomes containing unmodified histones have a higher interaction energy, whereas modifications like acetylation weaken internucleosomal interactions (58). Since the inactive chromatin examined here does not exhibit significant histone modifications (Supplementary Fig. 1 b), we have used a value from the upper range of the literature values (8 kT) to simulate this chromatin type. Conversely, the active region features many posttranslational histone modifications (Supplementary Fig. 1 a), and we thus used half the energy (4 kT) to compute the respective configurations.

### The nucleosome occupancy varies from cell to cell in active chromatin

Microscopic data shown so far are 2D data which underestimate the real 3D distances between the FISH spots since the cells are expected to be rotated randomly relative to the optical axis of the microscope. Only 3D single-cell microscopy allows the study of real distances between two spots on a single-cell level and to compare data between microscopy and simulation. Therefore, we performed 3D STED measurements which require careful correction for refractive index mismatch between immersion fluid of the objective lens and the embedding medium (see Materials and Methods).

The 3D STED measurements for the 5 kb AB interval in the active chromatin region revealed distances ranging from < 50 nm to 250 nm with a mean distance of 115 nm (n= 762, Fig. 4 a, data of all other segments Supplementary Fig. 2, statistical data in Supplementary Table 2). Remarkably, in active chromatin, elongated configurations can be found which results in a right-tailed distribution of the microscopic distance measurements. To understand this phenomenon better, we performed coarse-grained computer modelling of the nucleosome chain with the most probable nucleosome positions. We sampled a statistically relevant ensemble of independent 3D configurations in the active region applying our coarse-grained model that included elastic and electrostatic properties but excluded volume effects. In order to compare the simulated data with the microscopic data, the distances between the simulated sequence segments which correspond to those of the microscopic measurements were determined. In this way, a distance histogram was generated from the simulated data, which can be directly compared to the microscopic data (Fig. 4 b-g). The computed distribution was narrower, and the mean distance was about a standard deviation shorter than the microscopically measured distribution (Fig. 4 b).

**Fig. 4:**
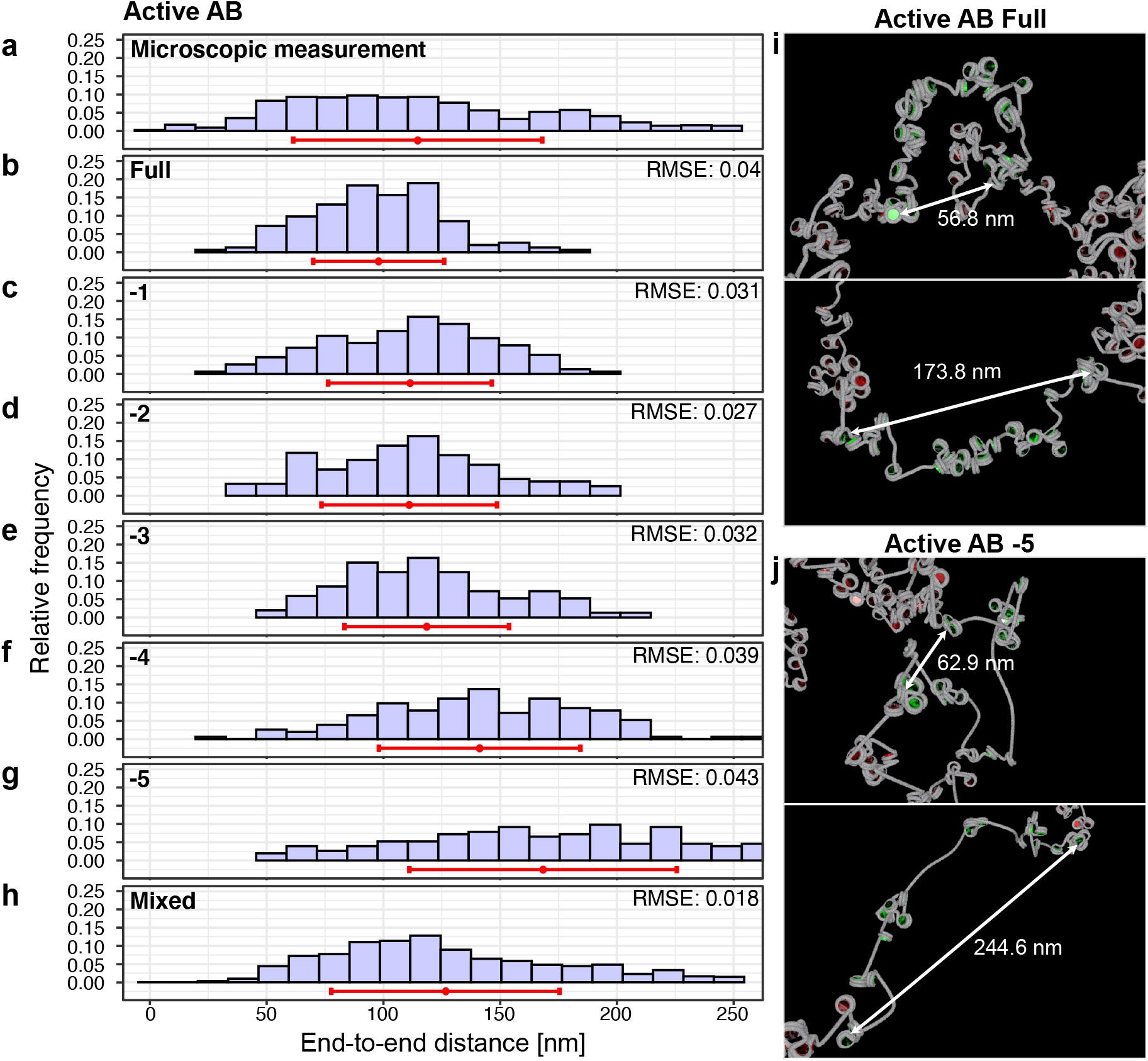
Distance distributions from microscopic experiments and from computer simulations of the active region. (a) 3D STED measurements of active AB result in a distance distribution ranging from < 50 nm to 250 nm with a mean of 115 nm ± 53 nm (n= 762 single-cell measurements from three independent replicates). (b-h) For computer simulations results are shown for the region active AB with all nucleosomes (Full) (b), with 1 to 5 nucleosomes replaced by naked DNA (c-g) and a combined plot (h). Mean value (red dot) and standard deviation (red line) are shown for each distribution. In the combined plot (h) the distributions have the weight 0.411, 0.000, 0.133, 0.000, 0.126, 0.330 (from all nucleosomes to −5 nucleosomes). (i-j) Example images of simulated chromatin fibers for active region AB (green nucleosomes) with all nucleosomes (i) and with 5 nucleosomes less (j) and the adjacent sequences (red nucleosomes). The upper image in (i) and (j) shows a configuration resulting in a short end-to-end distance indicated by a white arrow, the lower image depicts a large end-to-end distance. RMSE: root-mean-square error of simulated histogram bins in comparison to the measured data.

We hypothesized that under the physiological conditions of the microscopy experiment the number of bound nucleosomes varies from cell to cell. This hypothesis was tested by computer simulations, where the least probable nucleosomes were removed. To find the weakest bound nucleosomes, we analyzed the mean value from the occupancy data calculated by NucPosSimulator (weakest nucleosomes are indicated in Fig. 3 a). Next, we computed statistically relevant ensembles of 3D configurations by replacing the weakest nucleosome by naked DNA (−1, Fig. 4 c). The same was done by replacing two (Fig. 4 d), three (Fig. 4 e), four (Fig. 4 f) and five (Fig. 4 g) nucleosomes according to their rank order of binding strength. Indeed, a reduction of the total nucleosome number resulted in increasingly larger mean distances, but none of the individual distributions were comparable with the microscopically measured distribution. By applying a least squares fit, the different distance distributions were combined and resulted in a mixed distance histogram that mimics the microscopic data better than each of the underlying histograms as indicated by a reduction in the root-mean-square error (Fig. 4 h) (see Materials and Methods). Visualizations of simulated chromatin configurations show that both fibers with all nucleosomes and with a reduced nucleosome number (−5) can have short and long end-to-end distances (Fig. 4 i, j). These configurations show local accumulations of a few nucleosomes connected by stretches with low nucleosome occupancy. These structures are remarkably similar to recently published light and electron microscopic data of interphase chromatin (10, 65).

### Inactive region is compacted by various mechanisms

3D-STED distance histograms of the inactive region CD were compared with simulated data by the same strategy as above. The comparison showed that the computed mean distance was ~40 nm larger than the microscopically measured one when an attractive internucleosomal energy of 4 kT was used for the simulation (Fig. 5 a, b). As argued earlier, an increase of the interaction energy to 8 kT seems to be more realistic for simulating inactive chromatin. However, this approach delivered configurations with the mean value of the simulated distance distribution that are only a few nm shorter (Fig. 5 c). Obviously, additional mechanisms compact the inactive chromatin of the investigated region.

**Fig. 5:**
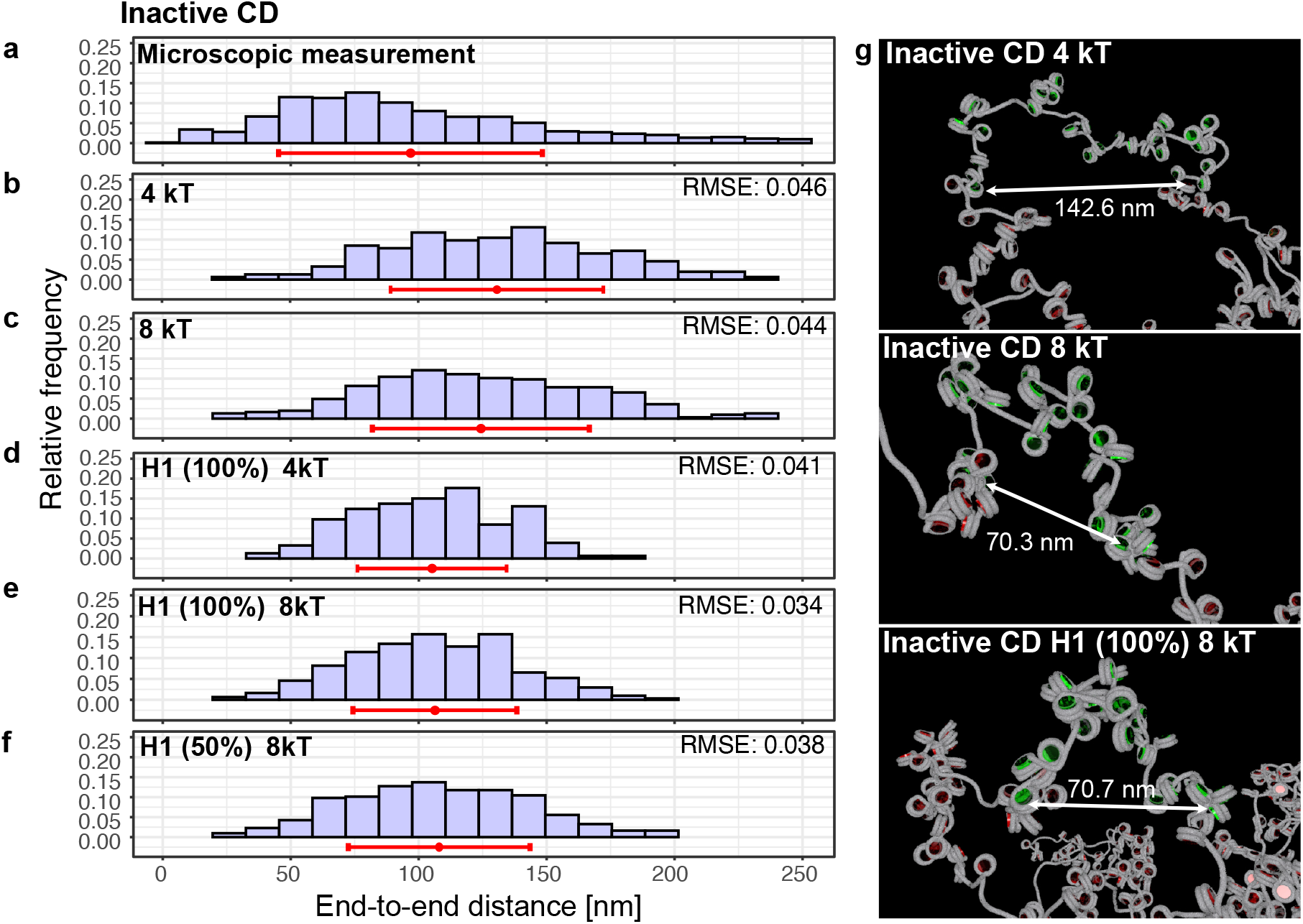
Distance distributions from microscopic experiments and from computer simulations of the inactive region. (a) 3D STED measurement of inactive CD results in a right-tailed distance distribution with the mass of the distribution towards shorter distances and a mean of 97 nm ± 52 nm (n= 1320 single-cell measurements from three independent replicates). (b-f) Computed distance distributions with different maximal internucleosomal interactions (4 kT (b, d) and 8 kT (c, e, f)), without (b, c) linker histone H1 or with (d, e) H1 (100% of nucleosomes occupied) and a random distribution of binding of 50% H1 (f). Mean value (red dot) and standard deviation (red line) are shown for each distribution. (g) Visualizations of simulated configurations.

ENCODE data show no pronounced histone modifications or repetitive DNA sequences in the inactive region CD, which makes chromatin compaction by binding of Polycomb group proteins or heterochromatinization unlikely. Therefore, other mechanisms must be considered, such as the binding of linker histone 1 (H1), which has long been known to have a chromatin-compacting effect (66). H1 is included in the computer model by different angles of the attached linker DNA at the nucleosomes (Kepper et al., 2008). These angles were derived by a systematic analysis of data from reconstituted fibers (67). It can be expected that details of the angles vary since the chicken linker histone H5, for example, causes different angles than human H1 (67). However, all variants of H1 lead to higher chromatin compaction.

In fact, simulations with a stochiometric H1 to nucleosome ratio of 1:1 led to more compact configurations. This effect is especially pronounced at 4 kT and weaker in simulations using a maximal internucleosomal attraction energy of 8 kT (Fig. 5 d, e). To explore the effects of different stoichiometry of H1 we performed computer simulations of a random 50% nucleosome binding (1:2). The width of the length distribution is widened only by a small amount (Fig. 5 f). Visualizations of exemplary simulated configurations are shown in Fig. 5 g. In summary, the efforts to shift the distance distribution to short values were partially successful. Larger distances as found in microscopic measurements might be caused by weakly bound nucleosomes as in active regions (cf. Fig 3 b).

## Discussion

By using high-throughput super-resolution microscopy, we studied the nanoscale organization of 5 kb chromatin segments which are located in active and inactive chromatin. The selected areas are prototypic for the respective chromatin class because patterns of prominent or absent DHS spreads over hundreds of kb around the selected region and can be found in more than 150 cell types. The nucleosomal repeat length is nearly the same for both regions and very close to the mean value of the whole chromosome. Considering the great similarity within the four active and four inactive intervals studied here, it can be assumed that the structural principles described apply to significant parts of the genome. In both active and inactive chromatin, 3D spatial distances between the endpoints of the 5 kb segments differ from cell to cell resulting in a broad right-tailed distance distribution with the mass of the distribution shifted more to shorter values in inactive chromatin. In contrast, simulations with different nucleosome occupancies, changed internucleosomal energies or deviations from stoichiometric H1 binding led to far narrower distance distributions. Therefore, the large width of the distance distribution seems to be a feature that is caused by the summation of cell-to-cell differences in the resulting histogram.

Unexpectedly, we found very elongated chromatin configurations with 5 kb exhibiting lengths of over 200 nm in both active and inactive chromatin. For comparison, a stretched beads-on-a-string chromatin fiber of 5 kb has a length of 243 nm (68). In simulations with our coarse-grained model elongated chromatin configurations are more probable if a number of nucleosomes is replaced by naked DNA. Therefore, it is important to investigate which nucleosomes have the weakest occupancy in our model. Indeed, 8 of the 10 most weakly bound nucleosomes in the active region are localized within DHSs (Supplementary Fig. 1 a), a result that is consistent with genome-wide measurements (41).

The perspective of cell-to-cell differences in nucleosome occupancy in active DNA is supported by different lines of evidence: (i) while at certain positions nucleosomes are positioned with high precision (69), nucleosome positions can vary substantially from cell to cell (40, 61), (ii) pioneer transcription factors and chromatin remodeling complexes can change nucleosome occupancy (70, 71), (iii) upregulation of genes is known to reduce the number of bound nucleosomes (39), (iv) transcription factors compete cooperatively with nucleosomes for access to DNA (72, 73), (v) regulatory elements are actuated in an all-or-none fashion by cooperative binding of transcriptional factors in place of a canonical nucleosome (41, 74).

In our simulation, DNA stretches without nucleosomes are handled as linker DNA with the respective elastic and electrostatic properties. However, in a physiological context evicted nucleosomes could be replaced by transcription factors as outlined above. Crystal structures show that in humans many TFs do not bend DNA, which also applies to members of the large family of TFs with a C2H2 zinc finger motif (75–77). This supports the conclusion that the replacement of nucleosomes by transcription factors may lead to an elongation of the DNA structure.

As described earlier, the microscopic measurements of inactive chromatin reveal a compaction that can be partially explained by an increase in the strength of internucleosomal interactions or by the additional introduction of the linker histone H1. Given the vast number of variables and mechanisms affecting nanoscale chromatin organization confidently identifying further mechanisms remains a challenge. We speculate that the density of the surrounding chromatin, which has not been taken into account in this study and by others, may play a role. Microscopic measurements show that the inactive region investigated here is expected to be embedded in a more compact chromatin environment (Supplementary Fig. 3). Indeed, preliminary modeling approaches reveal that the environment has a large influence on chromatin packing density. This mechanism might be particularly important for the inactive chromatin under investigation here, which lacks significant amounts of posttranslational histone modifications and therefore the measured compaction cannot be explained by heterochromatization by HP1 or Polycomb protein repression.

Microscopic data of the inactive region also exhibits elongated chromatin configurations (>200 nm) which can be best explained in our model by a reduced nucleosome occupancy varying from cell to cell. In fact, the data shown in Fig. 3 b support this hypothesis, as weakly bound nucleosomes also exist in inactive chromatin and could therefore explain not only the elongated configurations but also the wide distance distribution. Nucleosome eviction is only well studied in active chromatin regions, but the results shown here suggest that the phenomenon could also occur in inactive chromatin. The underlying mechanisms are most likely different from those in active chromatin. However, our interpretation of the data is that even inactive chromatin is subject to continuous reorganization.

An extensive body of literature (for review see (78)) on chromatin architecture focuses on the formation of chromatin loops bringing regulatory elements into close contact and thus regulating gene expression. Distances below which an enhancer is thought to activate a promotor range from less than 150 nm (53) to 300 nm (79). Here we show by high-throughput microscopy of human chromatin that in active regions more than 45% of the 5 kb endpoints approach to less than 100 nm whereas in inactive chromatin this is the case in more than 60% of the cells (value derived from data of Fig. 4 a and 5a). Interestingly, also inactive chromatin that does not have any active binding sites for regulatory factors shows significant incidents of end-to-end contacts. Apparently, thermodynamically driven spontaneous movements can bring regulatory elements into close contact with their promoters that are only a few kb distant from one another. Considering that 142.000 proximal enhancer-like elements can be found in the human genome at a distance of less than 2 kb (80), these spontaneous movements of chromatin could significantly influence gene regulation.

## Conclusion

Super-resolution microscopy of oligoFISH-labeled DNase I hypersensitive and insensitive interphase chromatin provides insights into its nanoscale organization. To this end, systematic measurements of physical distances between FISH labels separated by 5 kb of DNA were performed which revealed right-skewed distance distributions in both chromatin classes. In active chromatin, the labeled segments come closer than 100 nm in more than 45% of the investigated cells, a value that could be of importance for models of interactions between proximal promoters and enhancers. However, elongated configurations with distances larger than 200 nm were also found in active chromatin. Coarse-grained computer models demonstrated that elongated chromatin configurations are more likely when the nucleosome occupancy is reduced. The models reproduced microscopic data best when a cell-to-cell variability in nucleosome occupancy was assumed which is in agreement with a recent study using chromatin fiber sequencing. Microscopic measurements of inactive chromatin reveal more compact chromatin, with simulations of this region suggesting a combination of different compaction mechanisms. Surprisingly, elongated chromatin configurations also exist in more compact, DNase I-insensitive genomic regions.

## Material & Methods

### Key resources table

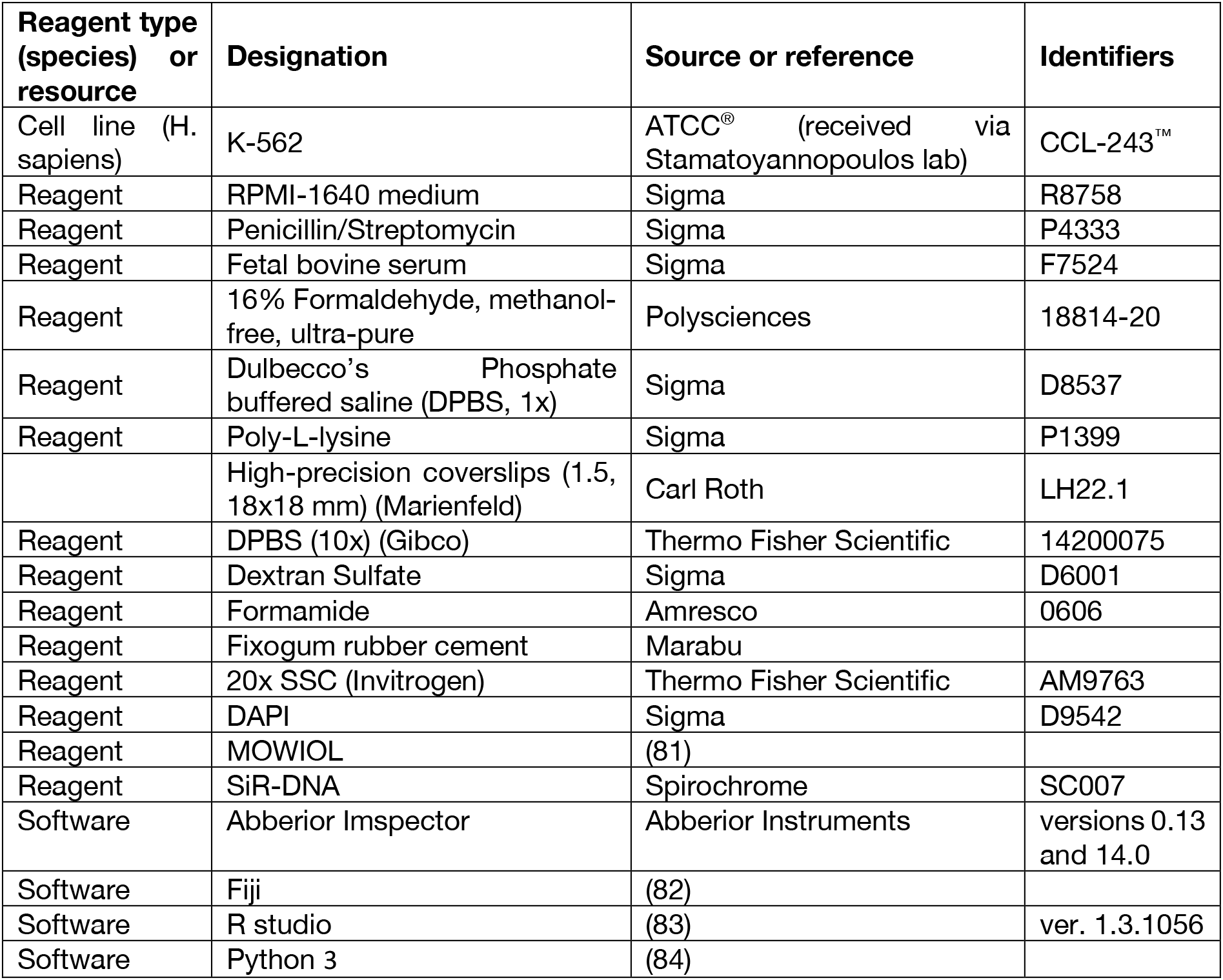

### Experimental model and subject details

#### Cell culture of K-562 cells

Human erythroleukemia K-562 cells were grown in RPMI-1640 medium (Sigma) supplemented with 10% FBS (Sigma) and 1% v/v penicillin/streptomycin (Sigma) in cell culture flasks. Cells were cultured at 37°C in 5% CO_2_ and regularly tested for mycoplasma contamination.

#### Selection criteria for regions used in this study

From the UCSC Genome browser (http://hgdownload.soe.ucsc.edu/goldenPath/hg38/database/), we downloaded the hg38 coordinates of centromeres, segmental duplications (“super dups”), and short tandem repeats (“RepeatMasker”). From the Gencode Genes project website, we downloaded version 37 of “basic” gene annotations for the 24 chromosomes in hg38 coordinates (ftp://ftp.ebi.ac.uk/pub/databases/gencode/Gencode_human/release_37/gencode.v37.basic.annotation.gtf.gz). From the Gencode file, we extracted the coordinates of gene bodies for genes annotated at level 1 or 2 for which at least one transcript is annotated at level 1 or 2 with a transcript support level of 1 or 2; this curated list of gene bodies comprises the “genic regions” in what follows. To assess chromatin accessibility across diverse cell and tissue types, we used the “Index” of (19) derived from DNase I hypersensitive regions called at FDR 0.1% in 733 diverse biosamples, and to assess accessibility in K-562 cells, we used regions called via the program hotspot2 at FDR 0.1% from the alignment file downloadable from https://www.encodeproject.org/files/ENCFF591TEM/.

To identify inactive regions, we took all genomic regions between successive elements in the Index, and the regions between the first and last Index elements on each chromosome and their respective ends of the chromosome. From these, we subtracted centromeric and genic regions, and retained all resulting regions with widths of at least 25 kb across which over 80% of the sites are uniquely mappable by 36mers and over 80% lie outside regions of segmental duplications. From these remaining regions, we chose the one that was overlapped the least by RepeatMasker elements. This region, chr11:55,810,260-55,840,940, stood out because less than 26.9% of it is overlapped by RepeatMasker elements; over 62% of each of the remaining candidate inactive regions are overlapped by RepeatMasker.

To identify active regions, we started by partitioning each chromosome into 50-kb segments, starting at the “left” end of each chromosome, and later repeating this 50-kb partitioning with an offset of 25kb into each chromosome. We restricted the 50-kb segments to those that are at least 50% overlapped by Index elements and overlapped <35% by RepeatMasker elements and not overlapped by any segmental duplications. We further restricted these to 50-kb regions fully containing an Index element present in 732 or 733 diverse biosamples and a strong (maximally-scoring) DNase I hypersensitive region in the K-562 biosample. We ranked the remaining 50-kb segments in descending order by the percentages by which they are overlapped by Index elements and considered their genic content and the degree to which they are overlapped by RepeatMasker elements. The region chr11:119,075,000-119,125,000 stood out for being spanned by a diverse set of genes.

In addition, the probe sets were spaced approximately 5 kb (midpoint to midpoint) from each other and, for the active region, were mostly placed on DHS peaks. The probe sets were designed to span 1.5 – 2 kb.

The probe sets were also transferred to the hg19 genome assembly by using the UCSC genome browser in order to be able to use the publicly available MNase-seq data set ENCSR000CXQ from the ENCODE project (see also Preparation and Simulation).

#### Oligonucleotide probes for STED microscopy

We tiled 30 non-overlapping oligonucleotides (40-mers) across each target region (1.5 - 2 kb), selected for uniqueness and a higher density than afforded by other published design tools optimized for whole genome coverage or chromosome walking (85–87). For STED microscopy, oligonucleotides were labeled with ATTO 594 or ATTO 647N (LGC Biosearch Technologies (Petaluma, CA)). Dye conjugation was carried out post-synthesis in a pool via an NHS-ester modification reaction. Working stocks of pools of 30 oligonucleotides covering the target regions had a total concentration of 10 µM and were diluted further for experiments. For a list of all oligo probes used in this study see Supplementary Table 4.

#### Sample preparation and fluorescence in situ hybridization

Hybridization was carried out as previously published with small adaptations (38). In brief, washed K-562 cells were resuspended in a small volume of PBS at a density of 1 million cells per ml and cell suspension was applied to poly-L-lysine (Sigma P1399) coated glass coverslips (1.5, 18×18 mm, Marienfeld). Cells were fixed using an osmotically balanced and methanol-free 4 % formaldehyde (Polysciences) solution which has previously been shown to not cause detectable nuclear shrinkage (59). The following washing and permeabilization steps were carried out according to Bintu et al (38). Coverslips were then inverted onto 8 µl of hybridization solution and sealed with rubber cement (Marabu). Slides were placed on a heat block set to 81 °C for 3 minutes. The samples were incubated at 37 °C over night (16 – 20 h). This protocol uses low hybridization temperatures and short hybridization times which has previously been shown to only minimally disrupt chromatin structure on the nanoscale by using electron and super-resolution microscopy (59, 88–90). Due to the directly labeled primary probes the protocol contains only washing steps on the second day. The samples were washed twice with 2x SSC for 15 minutes. Two 7-minute washes in 0.2x SSC/ 0.2 % Tween-20 were carried out on a heat block at 56 °C followed by one wash in 4x SSC/ 0.2 % Tween-20 at RT. DNA was counterstained with DAPI (100 μg/ml in 2x SSC), followed by two more washes in 2x SSC. Coverslips were mounted on microscopic slides with MOWIOL (2.5 % DABCO, pH 7.0) (modified from (81)), dried for 30 minutes and sealed with nail polish to preserve cell morphology and prevent shrinkage of cells.

#### Sample preparation for FISH and SiR-DNA staining

Samples were prepared the same way as for two color FISH. In this case only one probe pool (B for active and inactive region) with an ATTO 594 dye label was used for hybridization. Instead of DAPI counterstaining, the samples were stained in 2.5 μM SiR-DNA in 2x SSC for 1 h in a humid chamber. Subsequently, slides were washed two times with 2x SSC for 5 minutes. Coverslips were mounted on microscopic slides with MOWIOL (2.5 % DABCO, pH 7.0), dried for 30 min and sealed with nail polish.

#### STED microscopy for FISH two color imaging

Image acquisitions were carried out on a 3D STED microscope system from Abberior Instruments equipped with two pulsed excitation lasers (594, 0.3 mW and 640 nm, 1.2 mW), one pulsed depletion laser (775 nm, 1.2 W) and Avalanche photodiodes for detection. A 100x UPlanSApo 1.4 NA oil immersion objective (Olympus) was used for all acquisitions.

The STED hardware was controlled with Python scripts by using the specpy interface to the microscope control software Imspector (versions 0.13 and 14.0, Abberior Instruments). To find oligoFISH spot pairs confocal dual color 50 μm x 50 μm x 5 μm (for 2D and 3D acquisitions) volumes were acquired using 100 μm pinhole, 150 nm pixel size, 250 nm z-steps, 10 μs pixel dwell time, no line accumulation and excitation laser powers of 18.8% for 594 nm and 19.3 % for 640 nm. Confocal scans were investigated, points were detected with a Laplacian-of-Gaussian blob detector in both channels and nuclear regions exhibiting signals in both color channels no more than 5 pixels apart from one another were determined. At these points of interest, STED detail stacks (3 μm × 3μm × 1.4μm) were acquired. For 2D STED acquisitions, the spatial light modulator (SLM) was used to generate a 2D STED depletion pattern and stacks were acquired with 200 nm z steps, 7 planes, 20 nm pixel size, 10 μs pixel dwell time, 5x line accumulation, 100 μm pinhole, excitation laser power 53.5% for 594 nm, 53.5% for 640 nm and 29.6% for 775 nm depletion laser power. For 3D STED acquisitions, careful correction for refractive index mismatch between immersion fluid of the microscope objective and the cell is crucial. Therefore, immersion oil with a refractive index of 1.522 was used for 3D acquisitions. The SLM modulator was set to generate a 3D STED depletion pattern and stacks (3 μm × 3 μm × 1.5 μm) were imaged with 60 nm z steps, 25 planes, 45 nm pixel size, 10 μs pixel dwell time, 5x line accumulation, 100 μm pinhole, excitation laser power 53.5% for 594 nm, 53.5 % for 640 nm and 29.6% for 775 nm depletion laser power. The process was repeated for the next overview scan. The focus position was updated to the plane of maximum intensity in the previous overview image to allow for overnight imaging without focus loss. By moving the stage in x and y in a spiral pattern, overview scans followed by STED detail scans were acquired until a pre-set amount of time had passed.

#### STED microscopy for FISH and SiR-DNA co-imaging

Image acquisitions were carried out on a 3D STED microscope system from Abberior Instruments described above using a 100x UPlanSApo 1.4 NA oil immersion objective (Olympus). The STED hardware was controlled with Python scripts as described above. To find oligoFISH spots in 594 nm confocal dual color 50 μm × 50 μm × 7 μm volumes were acquired using 100 μm pinhole, 150 nm pixel size, 10 μs pixel dwell time, no line accumulation and excitation laser powers of 18.8 % for 594 nm and 19.3 % for 640 nm. Confocal scans were investigated, points were detected with a Laplacian-of-Gaussian blob detector in the 594 nm channel. At these points of interest, STED detail stacks (3 μm × 3 μm × 1.4 μm, 200 nm plane spacing) were acquired using the 594 nm laser for excitation. To get the surrounding SiR-DNA signal a 15 × 15 μm (30 nm pixel size, 1 plane) field of view was acquired around the same points of interest using the 640 nm laser. By moving the stage in x and y in a spiral pattern, overview scans followed by STED detail scans were acquired until a pre-set amount of time had passed.

#### STED microscopy image analysis for FISH spot distances

Though the automated data acquisition process produced large numbers of images, some of these were of insufficient quality for further analysis due to poor signal to noise ratio or spot detection only in one channel caused by premature bleaching or sample drift. Therefore, supervised machine learning was used as a quality control step to automatically classify STED stacks into ‘good’ or ‘bad’. An experienced scientist classified about more than two thousand sum projections of oligoFISH STED stacks as “analyzable data” or “not analyzable data”. Features extracted from the sum projections of his ground truth dataset were used to train a Random Forest classifier that could be used to automatically classify further acquisitions. All machine learning was done in Python using scikit-learn (ver. 0.19.1 or earlier). All acquired raw data including “good” and “bad” images can be found via https://osf.io/zjwxm/?view_only=1ee5eb2fe370422399e41d4e875b8771.

Detailed spot analysis was performed on the analyzable data to determine the coordinates of both FISH spots in their respective STED channels. The algorithm searched for the spot pair with the brightest signal and saved their subpixel coordinates for further statistical analysis. After a rough spot detection with a Laplacian-of-Gaussian blob detector, subpixel localization was performed by fitting a multidimensional Gaussian using the Levenberg-Marquardt algorithm. 3D coordinates were transformed into projected 2D coordinates by omitting the z coordinate. The code for handling the microscopy data and analysis is available at: https://bitbucket.org/davidhoerl/sted-oligofish-analysis

#### Chromatin environment of single FISH spots

To determine the relative chromatin compaction at the FISH spot, a maximum z-projection of the FISH stack was overlaid onto the single SiR-DNA plane (scaled with bilinear interpolation to match pixel sizes). In the resulting images, the spot position and nuclear outlines were annotated by hand. To reduce out-of-focus signal, a rolling-ball (radius=50px) background subtraction was performed on the SiR channel. For each image, the quantile of the SiR intensity at the FISH spot location with respect to all pixels in the nuclear annotation (smoothed with a Gaussian blur with sigma=1px) was determined. The results were visualized as boxplots and statistical significance of differences between inactive and active loci was assessed via a two-sided Wilcoxon rank sum test.

### 3D model

Since atomistic modelling of chains with many nucleosomes is not possible, coarse-grained models are widely used. We applied the simulation procedure as described in Müller et al. and we follow the description given there (50). Chromatin is modeled as a chain of segments, in which spherocylindrical units describing the nucleosomes are connected by cylindrical segments describing the linker DNA. Each segment *i* possesses a position and a local coordinate system consisting of three perpendicular unit vectors 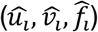 that describe its torsional orientation (Supplementary Fig. 4). Vector 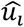 is parallel to the direction of the segment i.e. the vector 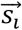 from its position to the position of the next segment. The position of the center of the nucleosome and its orientation is computed from the center of the nucleosome segment by the length d and 6 angles describing the relative orientation (Supplementary Fig. 5). Systems without linker histone and with linker histone differ by the set of angles (67).

The length of each individual linker DNA was computed from the positions of the nucleosomes in the studied region. The number of base pairs of a linker length is converted to nanometers by the factor of 0.34 nm/bp. Each linker DNA is modelled by at least 2 segments. If the linker length is larger than 20 nm the number of segments is calculated by rounding (linker length/10nm) up.

#### Simulation protocol

A Monte Carlo (MC) algorithm was utilized to create a statistically relevant set of configurations satisfying the Boltzmann distribution (91). In order to overcome local energy minima (51) we applied a replica exchange procedure introduced by Swendsen and Wang (92). Here, *M* replicas of the system were simulated with Metropolis Monte Carlo simultaneously, each at a different temperature *T*_*i*_. After a fixed number of MC simulation steps replicas with adjacent temperatures (*T*_*i*_, *T*_*i+1*_) the temperature is swapped with the probability:

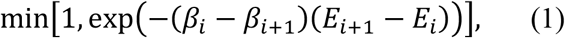

with *β*_*i*_ = 1/(*k*_*B*_*T*_*i*_), *k*_*B*_ being the Boltzmann constant and *E*_*i*_ the energy of the system i. Before the simulations the set of temperatures was determined utilizing a feedback-optimized approach (93). This algorithm optimizes the distribution of temperatures iteratively, such that the diffusion of replicas from the highest to the lowest temperature and vice versa is improved in each iteration. The procedure is more efficient when starting with a system that is pre-relaxed utilizing a simulated annealing approach (51).

#### Elastic energies

Elastic interactions are modelled by harmonic potentials. The strength constants of the interactions are named 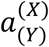 where *X* denotes the type of interaction (*s*=stretching, *b*=bending, *t*=torsion) and *Y* the interaction partners (DNA or nucleosome). The energy for stretching is calculated by:

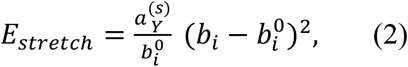

where *b*_*i*_ is the current length and 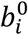 is the equilibrium length of the segment. The bending energy is given by:

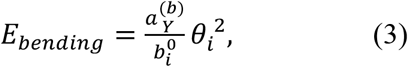

Where *θ*_*i*_ is calculated from 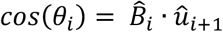 with 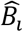 being the equilibrium direction of the next segment and 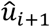 its actual direction. The torsional energy is computed as:

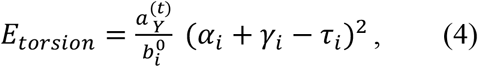

Where the angles *α*_*i*_, and *γ*_*i*_ are from the Euler-transformation (*α*_*i*_, *β*_*i*_, *γ*_*i*_) from the local coordinate system from segment *i* to segment *i+1*. The angle *τ*_*i*_ is the intrinsic twist (94).

#### Internucleosomal interaction

The internucleosomal interaction is described by a shifted 12-6 Lennard-Jones potential

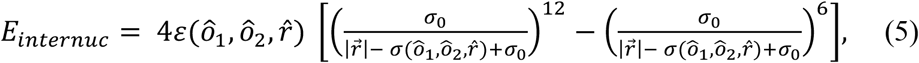

where *ô*_1_ and *ô*_2_ denote the orientation of the nucleosome and 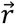 the distance between the centers of the nucleosomes. The shape of the nucleosome and the spatial dependency of the internucleosomal interaction strength is modelled by *ε* and *σ* depending of *ô*_1_, *ô*_2_ and 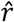. This is implemented by a series expansion in S-functions (95):

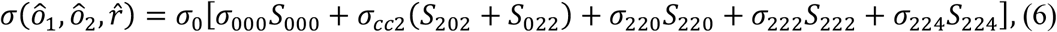

and

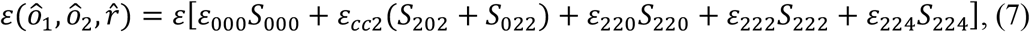

The expansion coefficients were chosen to match the spatial dimensions of the nucleosome and data from force spectroscopy experiments (51, 63, 67).

#### DNA-Nucleosome excluded volume

The volume of DNA segments is approximated by spheres. The minimal distance *d* between the center of DNA sphere and a spherocylinder describing the nucleosomes is computed. The excluded volume energies *E*_*DNA-Nuc*_, is described as the sum of the individual excluded volume energies *E′*_*DNA-Nuc*_, computed for DNA sphere and the volume of the nucleosome:

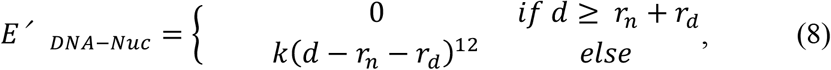

with *r*_*n*_ = (5.5/2) nm and *r*_*d*_ = 1.2 *nm*.

#### Electrostatic energy of linker DNA

A DNA segment is modelled by a chain c of charged spheres The GROMACS unit system was used which is based on nm, ps, K, electron charge (e) and atomic mass unit (u) (96).

The electrostatic energy of two spheres with charge *q*_1_ and *q*_2_ and radius a separated by a center-to-center distance *r* can be approximated by the electrostatic part of the Derjaguin-Landau-Verwey-Overbeek theory (97, 98) as

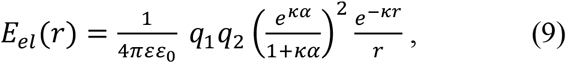

With *κ* being the inverse Debye length calculated by:

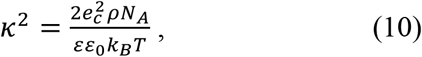

For the values listed in Supplementary Table 3 *κ* yields *κ* = 1.0387 *nm*^−1^ which corresponds to a Debye length of *λ*_*D*_ = *κ*^−1^ = 0.96 *nm*.

The charge of a DNA segment is given by *q* = *vd*, with *v* being the nominal line charge density (−2/0.34 *e*_*c*_, *nm*^−1^) and *d* the length of the DNA represented by the sphere. The line charge density *v* of the DNA must be adapted to the effective charge density *v* *

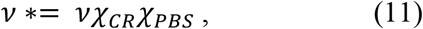

Where *χ*_*CR*_ is the charge adaptation factor and *χ*_*PBS*_ accounts for the geometry of subsequent overlapping beads and for deviations due to using an approximation instead of the exact Poisson-Boltzmann (PB) equation (99). Here, we use for *χ*_*CR*_ a value of 0.42 as derived in (99). The adaptation factor *χ*_*PBS*_ was determined by relating this potential to previous description as cylindrical segments (99).

#### Preparation and Simulation

For the preparation of the simulation data we first selected an appropriate human genome dataset (MNAse-seq of K-562 cells from the ENCODE project ENCSR000CXQ (15, 52)) in BigWig format (ENCFF000VNN). Next, we converted this file into the WIG-Format applying the BigWig2Wig-tool and finally in a BED format by a simple awk-script. Reads from chromosome 11 were extracted applying another simple UNIX-awk-script. In order to avoid false positive nucleosome positions blacklisted regions were filtered out (https://www.encodeproject.org/files/ENCFF001TDO/). Best nucleosome positions were determined with NucPosSimulator (61) generating a BED file containing the nucleosome positions and the occupancy, i.e. the number of read centers counted per base pair, smoothed with a Gaussian kernel and normalized. For identifying the least probable nucleosome the mean occupancy values of the 147 bp regions classified as nucleosomes by NucPosSimulator were determined and sorted. After removing the chosen number of nucleosomes with the smallest values, we generated a nucleosome chain with liker lengths as in the region and performed computer simulations (100). In order to incorporate effects of surrounding chromatin nucleosomes 20 kb were included at both sites of both investigated regions. The simulations were carried out on the linux cluster in Stralsund and the North German Supercomputing Alliance (HLRN) in Berlin.

#### Calculation of Nucleosome Repeat Length

The nucleosome repeat length (NRL) of whole chromosome 11 was determined analyzing the chr11 BED-file as described in the previous section. In a preparatory step nucleosome positions for the whole chromosome 11 were determined applying NucPosSimulator. From resulting sorted paired end nucleosome reads the repeat length between adjacent nucleosomes was calculated by subtracting the last base pair to the first base pair of the following nucleosome read. The average NRL a sliding window was calculated for a window size of 30000 bp. From this dataset windows with less than 3 nucleosomes e.g. in the centromere were removed applying filter-function from R package “dplyr” (filter(dataset(‘#Nucs’!=3))). The developed script (plotNRL.R) is published in a codeocean.com capsule (link).

#### Simulation Software

The software was developed in the Wedemann group in the last decades and used in many studies. It is written in C++ and was adapted for the use of shared-memory parallel architectures according to the OpenMP standard. The replica exchange algorithm was implemented for distributed memory architectures using Message Passing Interface (MPI). The simulation software was verified with an extensive set of unit tests and tests using simplified chain models which reproduced the expected analytical values. In addition, for visualization of chromatin configurations, a modular software was developed visualization of trajectories simulated by Monte Carlo procedures. The software cannot be made public at the moment, since it contains code under copyright by other parties.

#### Mixed histograms

A mixture histogram (Fig. 4h) was calculated by minimizing the squared differences between the bins of a histogram of the microscopically measured FISH spot distances (Fig. 4a) and a linear combination of the histograms of simulation results with varying nucleosome occupancies (Fig. 4b-g). Quadratic programming (via the quadprog package in R) was used to find a solution in which the contributions of the individual simulations are non-negative and sum to 1.

#### Statistics and reproducibility

No statistical method was used to predetermine sample size. Investigators were not blinded during the experiments and when assessing the outcome. For each experiment, data were collected from at least three independent biological replicates. 2D and 3D distance data were cut off at the maximum length of a theoretical beads-on-a-string fiber, since it is very unlikely that genomic regions are present in the nucleus more elongated than a fully stretched beads-on-a-string fiber. To calculate the length of a beads-on-a-string fiber the following formula was used: genomic length [bp] * 0.34 nm (size of one base) / 7 (68). For 5 kb genomic distances the cut-off for measured distances was at 250 nm. Plots in Fig. 2–5 and Supplementary Fig. 2 and 3 were generated using ggplot2 in R Studio (ver. 1.3.1056). Significance levels were always tested by a non-parametric two-sided Wilcoxon rank sum test and a Bonferroni-Holm correction was used to avoid errors through multiple testing when applicable. Data in Fig. 2–3 and Supplementary Fig. 3 are represented as boxplots where the middle line indicates the median, the lower and upper hinges correspond to the 25% and 75% quartiles, the upper whisker extends to the largest value no further than 1.5 × IQR (inter-quartile range) from the hinge and the lower whisker extends to the smallest value from the hinge at most 1.5 × IQR. The data acquisition, image processing and analysis was done in an unbiased way by automation.

## Availability of data and materials

Code used in this study for image processing and analysis can be found under https://bitbucket.org/davidhoerl/sted-oligofish-analysis. Code used in this study for processing of simulation data can be found as Code Ocean capsule via DOI 10.24433/CO.2730659.v1.

Datasets from microscopy and simulation generated and analyzed during the current study are available in the Open Science Framework repository under https://osf.io/zjwxm/?view_only=1ee5eb2fe370422399e41d4e875b8771.

Publicly available DNase-seq data sets with the ENCODE identifier ENCSR000EKS (UCSC Accession: wgEncodeEH000530 GEO accession: GSM816655) and ENCFF591TEM for K-562 (ENCSR000CXQ) cells were used and analyzed in the UCSC Genome Browser (http://genome-euro.ucsc.edu/). Publicly available MNase-seq data for K-562 cells (ENCSR000CXQ) were downloaded from https://www.encodeproject.org/ (15, 52). The data file with the identifier ENCFF000VNN was used. UCSC Accession: wgEncodeEH000921 GEO Accession: GSM920557

## Declarations

### Ethics approval and consent to participate

Not applicable

### Consent for publication

Not applicable

### Authors’ contributions

This study was conceived and supervised by HH, GW, JAS and HL. KB performed all microscopic experiments including sample preparation and STED imaging shown in Figures 1, 2, 4a, 5a and Supplementary Figures 2, 3. TR designed probes with help from ER and EH and provided input on sample preparation. KB analyzed and interpreted published ENCODE genome browser data with help from TR (Supplementary Figure 1). DH wrote scripts for microscope automation and image analysis. TZ performed computational modeling with input from GW (Figures 3, 4b-j, 5b-g, Supplementary Figures 4, 5). The manuscript was written by HH and GW with support from all authors.

## Acknowledgements

The authors acknowledge the North-German Supercomputing Alliance (HLRN) for providing HPC resources that have contributed to the research results reported in this paper. KB was supported by the International Max Planck Research School for Molecular Life Sciences (IMPRS-LS). Microscopic images were acquired at microscopes of the Center for Advacned Light Microscopy (CALM) at the LMU Munich. We thank Richard Sandstrom for help with Hi-C data.

## References

1. Trojer P, Reinberg D. Facultative heterochromatin: is there a distinctive molecular signature? Molecular cell. 2007;28(1):1–13.

2. Heitz E. Das Heterochromatin der Moose. Jahrb Wiss Bot. 1928;69:762–818.

3. Kornberg RD. Chromatin structure: a repeating unit of histones and DNA. Science. 1974;184(4139):868–71.

4. Olins AL, Olins DE. Spheroid chromatin units (ν bodies). Science. 1974;183(4122):330–2.

5. Luger K, Mader AW, Richmond RK, Sargent DF, Richmond TJ. Crystal structure of the nucleosome core particle at 2.8 A resolution. Nature. 1997;389(6648):251–60.

6. Mirny LA. The fractal globule as a model of chromatin architecture in the cell. Chromosome Res. 2011;19(1):37–51.

7. Maeshima K, Ide S, Babokhov M. Dynamic chromatin organization without the 30-nm fiber. Curr Opin Cell Biol. 2019;58:95–104.

8. Fussner E, Strauss M, Djuric U, Li R, Ahmed K, Hart M, et al. Open and closed domains in the mouse genome are configured as 10-nm chromatin fibres. EMBO Rep. 2012;13(11):992–6.

9. Konig P, Braunfeld MB, Sedat JW, Agard DA. The three-dimensional structure of in vitro reconstituted Xenopus laevis chromosomes by EM tomography. Chromosoma. 2007;116(4):349–72.

10. Ou HD, Phan S, Deerinck TJ, Thor A, Ellisman MH, O’Shea CC. ChromEMT: Visualizing 3D chromatin structure and compaction in interphase and mitotic cells. Science. 2017;357(6349).

11. Lakadamyali M, Cosma MP. Visualizing the genome in high resolution challenges our textbook understanding. Nature Methods. 2020.

12. Ernst J, Kellis M. Discovery and characterization of chromatin states for systematic annotation of the human genome. Nature biotechnology. 2010;28(8):817–25.

13. Ernst J, Kheradpour P, Mikkelsen TS, Shoresh N, Ward LD, Epstein CB, et al. Mapping and analysis of chromatin state dynamics in nine human cell types. Nature. 2011;473(7345):43–9.

14. Filion GJ, van Bemmel JG, Braunschweig U, Talhout W, Kind J, Ward LD, et al. Systematic protein location mapping reveals five principal chromatin types in Drosophila cells. Cell. 2010;143(2):212–24.

15. Consortium EP. An integrated encyclopedia of DNA elements in the human genome. Nature. 2012;489(7414):57–74.

16. Ram O, Goren A, Amit I, Shoresh N, Yosef N, Ernst J, et al. Combinatorial patterning of chromatin regulators uncovered by genome-wide location analysis in human cells. Cell. 2011;147(7):1628–39.

17. Hoffman MM, Buske OJ, Wang J, Weng Z, Bilmes JA, Noble WS. Unsupervised pattern discovery in human chromatin structure through genomic segmentation. Nature Methods. 2012;9(5):473–6.

18. Hoffman MM, Ernst J, Wilder SP, Kundaje A, Harris RS, Libbrecht M, et al. Integrative annotation of chromatin elements from ENCODE data. Nucleic acids research. 2013;41(2):827–41.

19. Meuleman W, Muratov A, Rynes E, Halow J, Lee K, Bates D, et al. Index and biological spectrum of human DNase I hypersensitive sites. Nature. 2020;584(7820):244–51.

20. Gross DS, Garrard WT. Nuclease hypersensitive sites in chromatin. Annual review of biochemistry. 1988;57(1):159–97.

21. Görisch SM, Wachsmuth M, Tóth KF, Lichter P, Rippe K. Histone acetylation increases chromatin accessibility. Journal of cell science. 2005;118(24):5825–34.

22. Moller J, Lequieu J, de Pablo JJ. The free energy landscape of internucleosome interactions and its relation to chromatin fiber structure. ACS central science. 2019;5(2):341–8.

23. Nozaki T, Imai R, Tanbo M, Nagashima R, Tamura S, Tani T, et al. Dynamic Organization of Chromatin Domains Revealed by Super-Resolution Live-Cell Imaging. Mol Cell. 2017;67(2):282–93 e7.

24. Zhang R, Erler J, Langowski J. Histone acetylation regulates chromatin accessibility: role of H4K16 in inter-nucleosome interaction. Biophysical journal. 2017;112(3):450–9.

25. Otterstrom J, Castells-Garcia A, Vicario C, Gomez-Garcia PA, Cosma MP, Lakadamyali M. Super-resolution microscopy reveals how histone tail acetylation affects DNA compaction within nucleosomes in vivo. Nucleic acids research. 2019;47(16):8470–84.

26. Allshire RC, Madhani HD. Ten principles of heterochromatin formation and function. Nature Reviews Molecular Cell Biology. 2018;19(4):229.

27. Heun P, Laroche T, Shimada K, Furrer P, Gasser SM. Chromosome dynamics in the yeast interphase nucleus. Science. 2001;294(5549):2181–6.

28. Marshall W, Straight A, Marko J, Swedlow J, Dernburg A, Belmont A, et al. Interphase chromosomes undergo constrained diffusional motion in living cells. Current Biology. 1997;7(12):930–9.

29. Levi V, Ruan Q, Plutz M, Belmont AS, Gratton E. Chromatin dynamics in interphase cells revealed by tracking in a two-photon excitation microscope. Biophysical journal. 2005;89(6):4275–85.

30. Hajjoul H, Mathon J, Ranchon H, Goiffon I, Mozziconacci J, Albert B, et al. High-throughput chromatin motion tracking in living yeast reveals the flexibility of the fiber throughout the genome. Genome research. 2013;23(11):1829–38.

31. Lucas JS, Zhang Y, Dudko OK, Murre C. 3D trajectories adopted by coding and regulatory DNA elements: first-passage times for genomic interactions. Cell. 2014;158(2):339–52.

32. Germier T, Kocanova S, Walther N, Bancaud A, Shaban HA, Sellou H, et al. Real-time imaging of a single gene reveals transcription-initiated local confinement. Biophysical journal. 2017;113(7):1383–94.

33. Chen B, Gilbert LA, Cimini BA, Schnitzbauer J, Zhang W, Li G-W, et al. Dynamic imaging of genomic loci in living human cells by an optimized CRISPR/Cas system. Cell. 2013;155(7):1479–91.

34. Gu B, Swigut T, Spencley A, Bauer MR, Chung M, Meyer T, et al. Transcription-coupled changes in nuclear mobility of mammalian cis-regulatory elements. Science. 2018.

35. Ma H, Tu LC, Chung YC, Naseri A, Grunwald D, Zhang S, et al. Cell cycle- and genomic distance-dependent dynamics of a discrete chromosomal region. J Cell Biol. 2019.

36. Shaban HA, Barth R, Bystricky K. Formation of correlated chromatin domains at nanoscale dynamic resolution during transcription. Nucleic acids research. 2018;46(13):e77–e.

37. Zidovska A, Weitz DA, Mitchison TJ. Micron-scale coherence in interphase chromatin dynamics. Proceedings of the National Academy of Sciences. 2013;110(39):15555–60.

38. Bintu B, Mateo LJ, Su J-H., Sinnott-Armstrong NA, Parker M, Kinrot S, et al. Super-resolution chromatin tracing reveals domains and cooperative interactions in single cells. Science. 2018;362(6413):eaau1783.

39. Diermeier S, Kolovos P, Heizinger L, Schwartz U, Georgomanolis T, Zirkel A, et al. TNFα signalling primes chromatin for NF-κB binding and induces rapid and widespread nucleosome repositioning. Genome biology. 2014;15(12):536.

40. Lai B, Gao W, Cui K, Xie W, Tang Q, Jin W, et al. Principles of nucleosome organization revealed by single-cell micrococcal nuclease sequencing. Nature. 2018;562(7726):281–5.

41. Stergachis AB, Debo BM, Haugen E, Churchman LS, Stamatoyannopoulos JA. Single-molecule regulatory architectures captured by chromatin fiber sequencing. Science. 2020;368(6498):1449–54.

42. Becker PB, Workman JL. Nucleosome remodeling and epigenetics. Cold Spring Harbor Perspectives in Biology. 2013;5(9):a017905.

43. Hargreaves DC, Crabtree GR. ATP-dependent chromatin remodeling: genetics, genomics and mechanisms. Cell research. 2011;21(3):396–420.

44. Dultz E, Mancini R, Polles G, Vallotton P, Alber F, Weis K. Quantitative imaging of chromatin decompaction in living cells. Mol Biol Cell. 2018:mbcE17110648.

45. Parmar JJ, Padinhateeri R. Nucleosome positioning and chromatin organization. Current Opinion in Structural Biology. 2020;64:111–8.

46. Collepardo-Guevara R, Schlick T. Chromatin fiber polymorphism triggered by variations of DNA linker lengths. Proceedings of the National Academy of Sciences. 2014;111(22):8061–6.

47. Clauvelin N, Lo P, Kulaeva O, Nizovtseva E, Diaz-Montes J, Zola J, et al. Nucleosome positioning and composition modulate in silico chromatin flexibility. Journal of Physics: Condensed Matter. 2015;27(6):064112.

48. Nordenskiöld L, Lyubartsev AP, Korolev N. Coarse-grained modeling of nucleosomes and chromatin: CRC Press; 2017.

49. Kepper N, Foethke D, Stehr R, Wedemann G, Rippe K. Nucleosome geometry and internucleosomal interactions control the chromatin fiber conformation. Biophys J. 2008;95(8):3692–705.

50. Muller O, Kepper N, Schopflin R, Ettig R, Rippe K, Wedemann G. Changing chromatin fiber conformation by nucleosome repositioning. Biophys J. 2014;107(9):2141–50.

51. Stehr R, Kepper N, Rippe K, Wedemann G. The effect of internucleosomal interaction on folding of the chromatin fiber. Biophys J. 2008;95(8):3677–91.

52. Davis CA, Hitz BC, Sloan CA, Chan ET, Davidson JM, Gabdank I, et al. The Encyclopedia of DNA elements (ENCODE): data portal update. Nucleic Acids Res. 2018;46(D1):D794–d801.

53. Mateo LJ, Murphy SE, Hafner A, Cinquini IS, Walker CA, Boettiger AN. Visualizing DNA folding and RNA in embryos at single-cell resolution. Nature. 2019;568(7750):49–54.

54. Sahl SJ, Hell SW, Jakobs S. Fluorescence nanoscopy in cell biology. Nature reviews Molecular cell biology. 2017;18(11):685.

55. Göttfert F, Wurm CA, Mueller V, Berning S, Cordes VC, Honigmann A, et al. Coaligned dual-channel STED nanoscopy and molecular diffusion analysis at 20 nm resolution. Biophysical journal. 2013;105(1):L01–L3.

56. Finn EH, Pegoraro G, Brandao HB, Valton AL, Oomen ME, Dekker J, et al. Extensive Heterogeneity and Intrinsic Variation in Spatial Genome Organization. Cell. 2019;176(6):1502–15 e10.

57. Ashwin SS, Maeshima K, Sasai M. Heterogeneous fluid-like movements of chromatin and their implications to transcription. Biophys Rev. 2020;12(2):461–8.

58. Funke JJ, Ketterer P, Lieleg C, Schunter S, Korber P, Dietz H. Uncovering the forces between nucleosomes using DNA origami. Science advances. 2016;2(11):e1600974.

59. Boettiger AN, Bintu B, Moffitt JR, Wang S, Beliveau BJ, Fudenberg G, et al. Super-resolution imaging reveals distinct chromatin folding for different epigenetic states. Nature. 2016;529(7586):418–22.

60. Cui K, Zhao K. Genome-wide approaches to determining nucleosome occupancy in metazoans using MNase-Seq. Methods Mol Biol. 2012;833:413–9.

61. Schopflin R, Teif VB, Muller O, Weinberg C, Rippe K, Wedemann G. Modeling nucleosome position distributions from experimental nucleosome positioning maps. Bioinformatics. 2013;29(19):2380–6.

62. Mangenot S, Leforestier A, Vachette P, Durand D, Livolant F. Salt-induced conformation and interaction changes of nucleosome core particles. Biophysical journal. 2002;82(1):345–56.

63. Kepper N, Ettig R, Stehr R, Marnach S, Wedemann G, Rippe K. Force spectroscopy of chromatin fibers: extracting energetics and structural information from Monte Carlo simulations. Biopolymers. 2011;95(7):435–47.

64. Norouzi D, Zhurkin VB. Dynamics of chromatin fibers: comparison of monte carlo simulations with force spectroscopy. Biophysical journal. 2018;115(9):1644–55.

65. Ricci MA, Manzo C, Garcia-Parajo MF, Lakadamyali M, Cosma MP. Chromatin fibers are formed by heterogeneous groups of nucleosomes in vivo. Cell. 2015;160(6):1145–58.

66. Van Holde KE. Chromatin. Heidelberg: Springer Science & Business Media; 1989.

67. Stehr R, Schopflin R, Ettig R, Kepper N, Rippe K, Wedemann G. Exploring the conformational space of chromatin fibers and their stability by numerical dynamic phase diagrams. Biophys J. 2010;98(6):1028–37.

68. Carlson RD, Olins DE. Chromatin model calculations: Arrays of spherical ν bodies. Nucleic Acids Research. 1976;3(1):89–100.

69. Baldi S, Korber P, Becker PB. Beads on a string—nucleosome array arrangements and folding of the chromatin fiber. Nature structural & molecular biology. 2020;27(2):109–18.

70. Zaret KS. Pioneer Transcription Factors Initiating Gene Network Changes. Annual Review of Genetics. 2020;54.

71. Bartholomew B. Regulating the Chromatin Landscape: Structural and Mechanistic Perspectives. Annual Review of Biochemistry. 2014;83(1):671–96.

72. Svaren J, Klebanow E, Sealy L, Chalkley R. Analysis of the competition between nucleosome formation and transcription factor binding. Journal of Biological Chemistry. 1994;269(12):9335–44.

73. Mirny LA. Nucleosome-mediated cooperativity between transcription factors. Proceedings of the National Academy of Sciences. 2010;107(52):22534–9.

74. Thurman RE, Rynes E, Humbert R, Vierstra J, Maurano MT, Haugen E, et al. The accessible chromatin landscape of the human genome. Nature. 2012;489(7414):75–82.

75. Panne D, Maniatis T, Harrison SC. An atomic model of the interferon-β enhanceosome. Cell. 2007;129(6):1111–23.

76. Kim CA, Berg JM. A 2.2 Å resolution crystal structure of a designed zinc finger protein bound to DNA. Nature structural biology. 1996;3(11):940–5.

77. Pavletich NP, Pabo CO. Zinc finger-DNA recognition: crystal structure of a Zif268-DNA complex at 2.1 A. Science. 1991;252(5007):809–17.

78. Schoenfelder S, Fraser P. Long-range enhancer-promoter contacts in gene expression control. Nat Rev Genet. 2019;20(8):437–55.

79. Chen H, Levo M, Barinov L, Fujioka M, Jaynes JB, Gregor T. Dynamic interplay between enhancer–promoter topology and gene activity. Nature genetics. 2018;50(9):1296–303.

80. Moore JE, Purcaro MJ, Pratt HE, Epstein CB, Shoresh N, Adrian J, et al. Expanded encyclopaedias of DNA elements in the human and mouse genomes. Nature. 2020;583(7818):699–710.

81. Wurm CA, Neumann D, Schmidt R, Egner A, Jakobs S. Sample preparation for STED microscopy. Live cell imaging: Springer; 2010. p. 185–99.

82. Schindelin J, Arganda-Carreras I, Frise E, Kaynig V, Longair M, Pietzsch T, et al. Fiji: an open-source platform for biological-image analysis. Nat Methods. 2012;9(7):676–82.

83. RStudioTeam. RStudio: Integrated Development for R. RStudio, PBC, Boston, MA. 2020.

84. Van Rossum G, Drake, F. L. Python 3 Reference Manual. Scotts Valley, CA: Create Space. 2009.

85. Beliveau BJ, Kishi JY, Nir G, Sasaki HM, Saka SK, Nguyen SC, et al. OligoMiner provides a rapid, flexible environment for the design of genome-scale oligonucleotide in situ hybridization probes. Proc Natl Acad Sci U S A. 2018;115(10):E2183–E92.

86. Gelali E, Girelli G, Matsumoto M, Wernersson E, Custodio J, Mota A, et al. iFISH is a publically available resource enabling versatile DNA FISH to study genome architecture. Nature Communications. 2019;10(1):1636.

87. Beliveau BJ, Joyce EF, Apostolopoulos N, Yilmaz F, Fonseka CY, McCole RB, et al. Versatile design and synthesis platform for visualizing genomes with Oligopaint FISH probes. Proc Natl Acad Sci U S A. 2012;109(52):21301–6.

88. Markaki Y, Smeets D, Fiedler S, Schmid VJ, Schermelleh L, Cremer T, et al. The potential of 3D-FISH and super-resolution structured illumination microscopy for studies of 3D nuclear architecture: 3D structured illumination microscopy of defined chromosomal structures visualized by 3D (immuno)-FISH opens new perspectives for studies of nuclear architecture. Bioessays. 2012;34(5):412–26.

89. Solovei I, Cavallo A, Schermelleh L, Jaunin F, Scasselati C, Cmarko D, et al. Spatial preservation of nuclear chromatin architecture during three-dimensional fluorescence in situ hybridization (3D-FISH). Exp Cell Res. 2002;276(1):10–23.

90. Branco MR, Pombo A. Intermingling of chromosome territories in interphase suggests role in translocations and transcription-dependent associations. PLoS Biol. 2006;4(5):e138.

91. Metropolis N, Rosenbluth AW, Rosenbluth MN, Teller AH, Teller E. Equation of state calculations by fast computing machines. The journal of chemical physics. 1953;21(6):1087–92.

92. Swendsen RH, Wang JS. Replica Monte Carlo simulation of spin glasses. Phys Rev Lett. 1986;57(21):2607–9.

93. Katzgraber HG, Trebst S, Huse DA, Troyer M. Feedback-optimized parallel tempering Monte Carlo. Journal of Statistical Mechanics: Theory and Experiment. 2006;2006(03):P03018.

94. Klenin K, Merlitz H, Langowski J. A Brownian dynamics program for the simulation of linear and circular DNA and other wormlike chain polyelectrolytes. Biophys J. 1998;74(2 Pt 1):780–8.

95. Zewdie H. Computer simulation studies of liquid crystals: A new Corner potential for cylindrically symmetric particles. The Journal of chemical physics. 1998;108(5):2117–33.

96. Hess B, Kutzner C, van der Spoel D, Lindahl E. GROMACS 4: Algorithms for Highly Efficient, Load-Balanced, and Scalable Molecular Simulation. J Chem Theory Comput. 2008;4(3):435–47.

97. Levin Y. Electrostatic correlations: from plasma to biology. Reports on progress in physics. 2002;65(11):1577.

98. Walker DA, Kowalczyk B, de la Cruz MO, Grzybowski BA. Electrostatics at the nanoscale. Nanoscale. 2011;3(4):1316–44.

99. Maffeo C, Schopflin R, Brutzer H, Stehr R, Aksimentiev A, Wedemann G, et al. DNA-DNA interactions in tight supercoils are described by a small effective charge density. Phys Rev Lett. 2010;105(15):158101.

100. Mörl M-C., Zülske T, Schöpflin R, Wedemann G. Data formats for modelling the spatial structure of chromatin based on experimental positions of nucleosomes. AIMS Biophysics. 2019;6(3):83.

