## Supplementary Material for "Differences in nanoscale organization of DNase I hypersensitive and insensitive chromatin in single human cells"

**Brandstetter et al.**

**Supplementary Figures 1 – 5**
**Supplementary Table 1 – 4**

### Supplementary figures with explanation

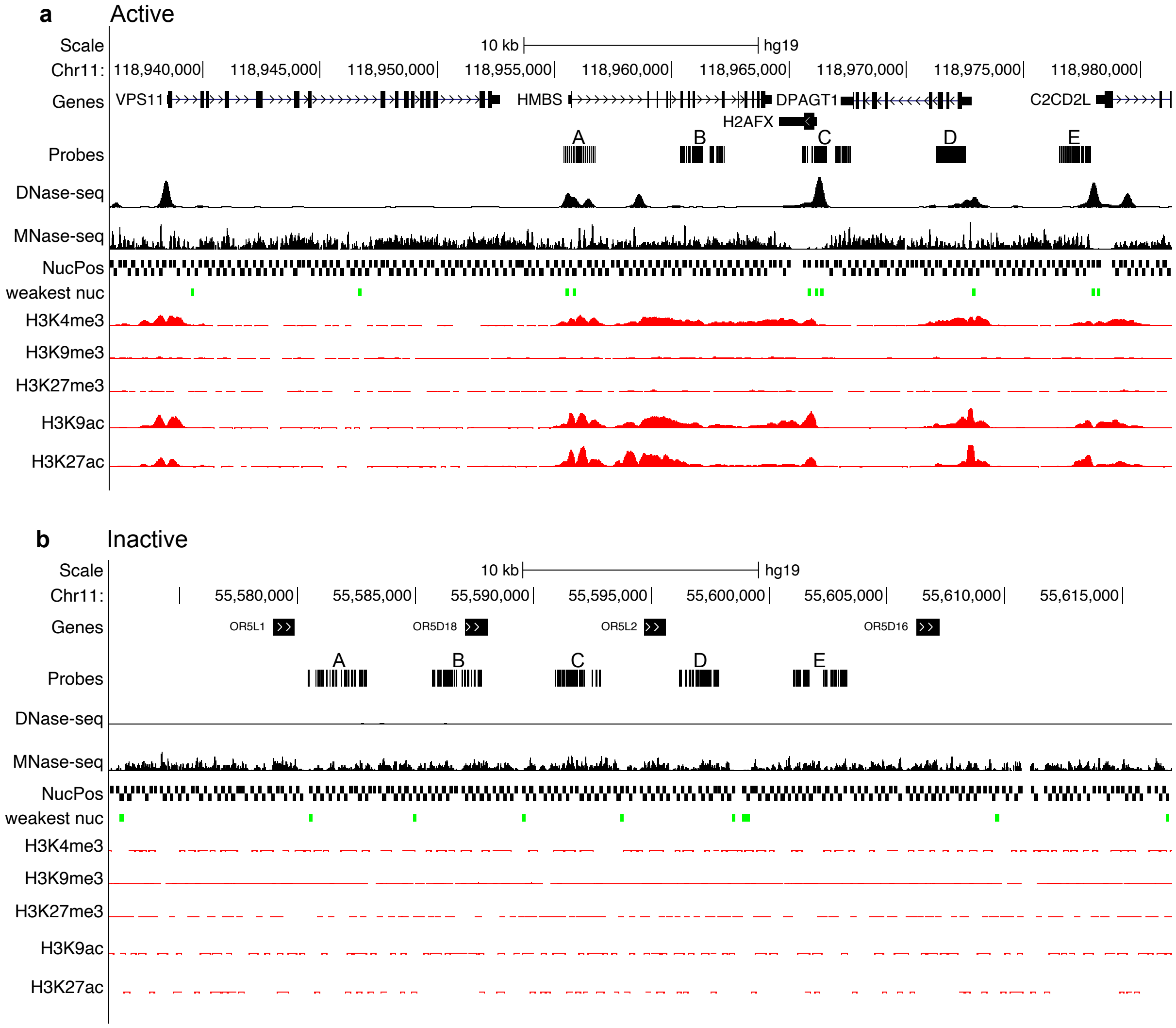

**Supplementary Figure 1:** Active (a) and inactive (b) genomic regions with genes, probe sets (A-E), DNase-seq, MNase-seq, nucleosome positions from NucPosSimulator, the weakest nucleosomes calculated by NucPosSimulator, H3K4me3, H3K9me3, H3K27me3, H3K9ac, H3K27ac. Tracks show that inactive region has almost no histone modifications while the active region contains active marks like H3K4me3, H3K9ac and H3K27ac. Notably, most of the weakest nucleosomes for the active region are located at DNase I hypersensitive sites.

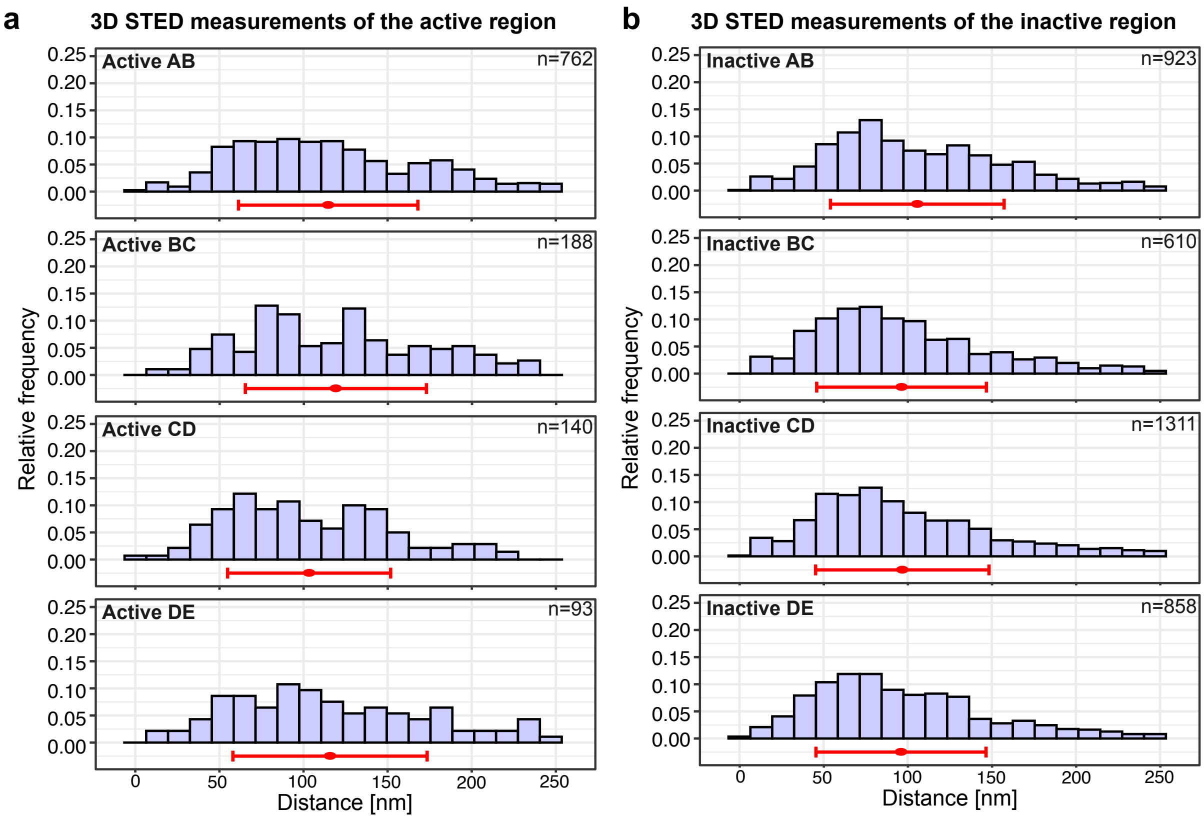

**Supplementary Figure 2:** Distance histograms from 3D STED measurements for all four intervals (AB, BC, CD, DE) in active (a) and inactive (b). The mean for each histogram is indicated by the red dot and the standard deviation by the red line. N-numbers can be found next to the respective histogram (for statistical data, see Supplementary Table 2).

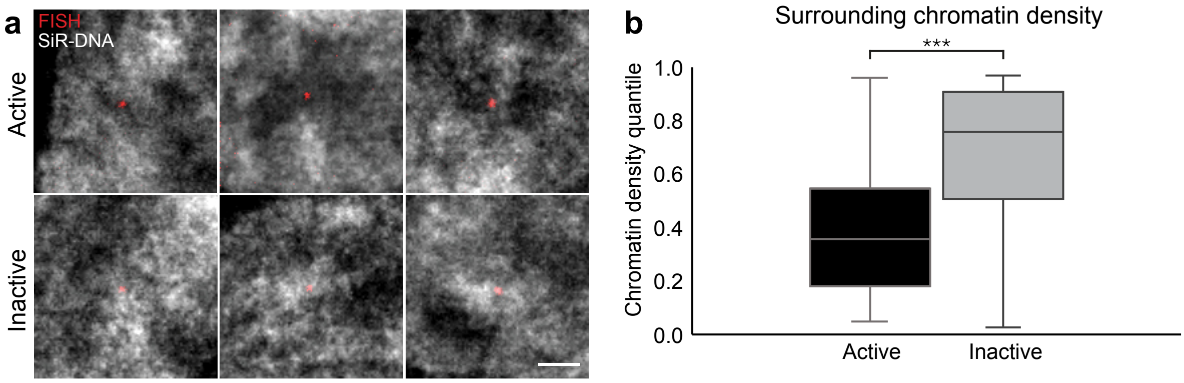

**Supplementary Figure 3:** Chromatin environment of the active and inactive region. (a) Representative images for the active (upper row) and inactive (lower row) region labeled with one FISH probe set (red) and chromatin labeled with SiR-DNA (gray). Scale bar = 1 µm. (b) Chromatin density quantile for active (black, n=43) and inactive (gray, n=38) differ significantly (two-sided Wilcoxon rank sum test, $p = 9.890 * 10^{-6}$). Inactive region is embedded in higher density chromatin, while active chromatin is surrounded by lower density chromatin.

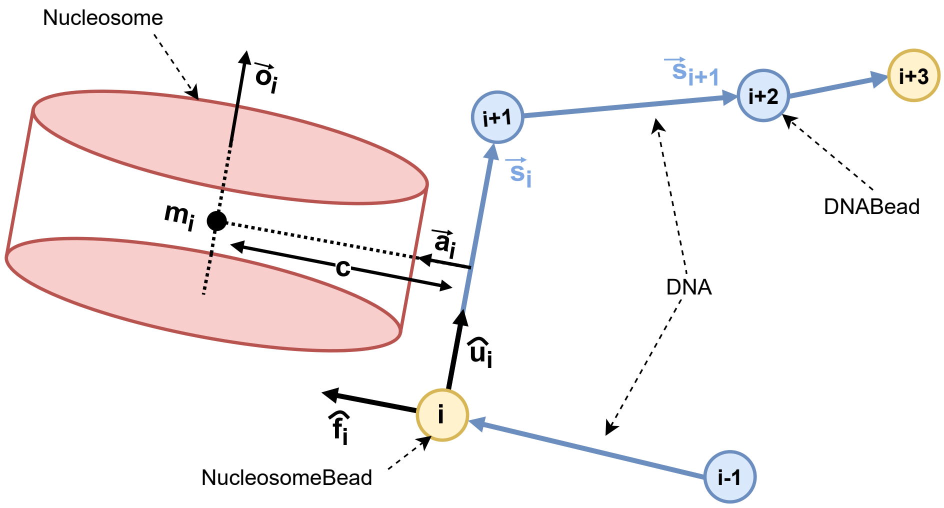

**Supplementary Figure 4:** Model of a nucleosome chain. *i* represents the position of the bead in the chain, yellow circles indicate nucleosome bead positions, and blue circles indicate DNA bead positions. The nucleosome is represented by a red cylinder. The segment vector $\vec{s_i}$ points from one bead to the next bead. A local coordination system $(\hat{u}_i, \hat{v}_i, \hat{f}_i)$ (not shown) describes the orientation of a bead. Vector $\vec{a_i}$ describes the direction from the center of the segment to the nucleosome center $m_i$, c is its length, and vector $\vec{o_i}$ describes the orientation of the nucleosome. Vector $\vec{a_i}$ is defined by two rotations of vector $\hat{v}_i$ (i) around $\vec{u_i}$ by the angle $\varepsilon$ (not shown), (ii) around vector $\hat{f}_i$ by the angle $\phi$ (not shown).

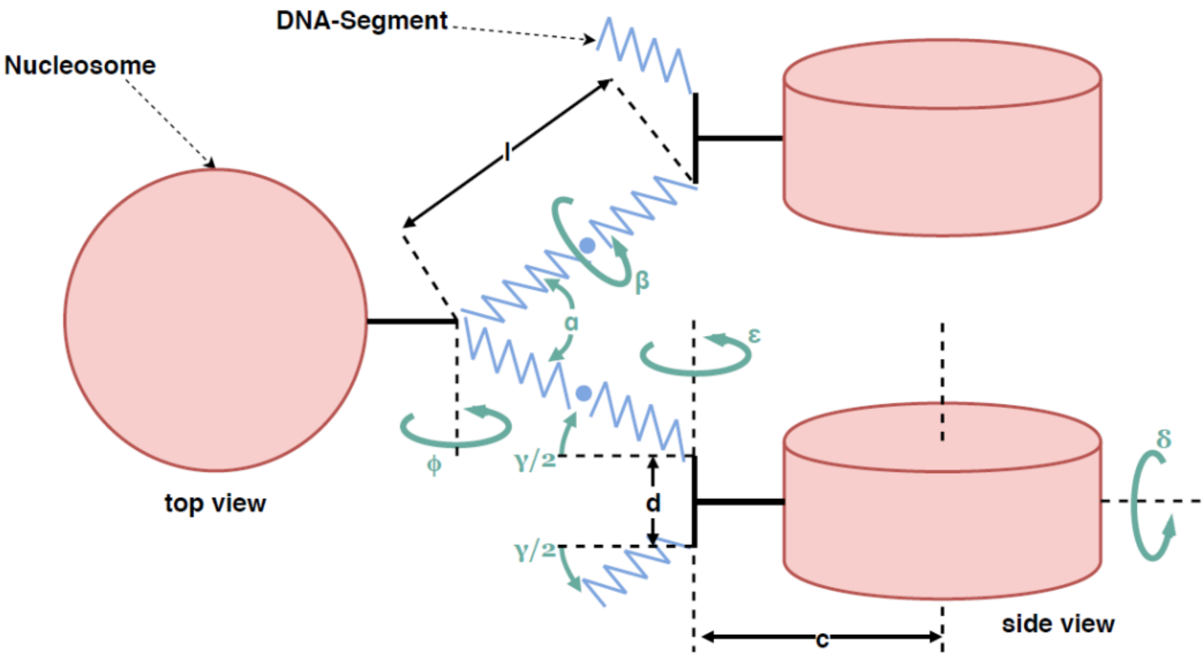

**Supplementary Figure 5**: The relative orientation of the nucleosome is described by the angles $\alpha, \delta, \varepsilon, \gamma, \phi$ (modified from (Rippe et al., 2012)). $\beta$ is the torsional orientation of subsequent nucleosomes. $l$ is the length of the DNA modeling the linker DNA, d the distance between the entry and the exit point of the linker DNA at the nucleosome, c the distance between the center of the nucleosome segment and the center of the oblate sphero-cylinder modeling of the nucleosome.

#### Supplementary tables with explanation

**Supplementary Table 1: Statistical data for 2D STED datasets**
Test: Wilcoxon rank sum test with continuity correction
Correction method: Bonferroni holm correction for multiple testing
Software: R studio

p-values for Active AB – BC – CD – DE

| | **Active AB** | **Active BC** | **Active CD** |
|---|---|---|---|
| **Active BC** | 0.03220 | - | - |
| **Active CD** | 0.02706 | 0.80123 | - |
| **Active DE** | 0.18970 | 0.00053 | 0.00053 |

p-values for Inactive AB – BC – CD – DE

| | **Inactive AB** | **Inactive BC** | **Inactive CD** |
|---|---|---|---|
| **Inactive BC** | $3.3 * 10^{-7}$ | - | - |
| **Inactive CD** | $5.8 * 10^{-9}$ | 0.451 | - |
| **Inactive DE** | $1.2 * 10^{-12}$ | 0.071 | 0.451 |

p-value for all active vs. all inactive

| | **Active** |
|---|---|
| **Inactive** | $<2 * 10^{-16}$ |

**Supplementary Table 2: Statistical data for 3D STED datasets**
Test: Wilcoxon rank sum test with continuity correction
Correction method: Bonferroni holm correction for multiple testing
Software: R studio
Significant: $p < 0.05$

p-values for Active AB – BC – CD – DE

| | **Active AB** | **Active BC** | **Active CD** |
|---|---|---|---|
| **Active BC** | 0.895 | - | - |
| **Active CD** | 0.138 | 0.076 | - |
| **Active DE** | 1.000 | 1.000 | 0.525 |

p-values for Inactive AB – BC – CD – DE

| | **Inactive AB** | **Inactive BC** | **Inactive CD** |
|---|---|---|---|
| **Inactive BC** | $8 * 10^{-4}$ | - | - |
| **Inactive CD** | $8.3 * 10^{-5}$ | 1.000 | - |
| **Inactive DE** | $9.4 * 10^{-5}$ | 1.000 | 1.000 |

p-value for all active vs. all inactive

| | **Active** |
|---|---|
| **Inactive** | $<2 * 10^{-16}$ |

**Supplementary Table 3: Simulation Parameters and Constants**

| | | |
|---|---|---|
| $e_c$ | $1.602 \times 10^{-19}$ C | Electric charge unit |
| $v$ | $-2/0.34\ e_c$ nm$^{-1}$ | Line charge density of DNA |
| $\rho$ | $0.1 \times 10^{24}$ mol nm$^{-3}$ | Molarity of the monovalent solution |
| $N_A$ | $6.022 \times 10^{23}$ mol$^{-1}$ | Avogadro constant |
| $\varepsilon$ | 80 | Value for the dielectric value in the solution |
| $\varepsilon_0$ | $(4\pi f)^{-1}$ | Dielectric constant |
| $f$ | 138.935 kJ nm mol$^{-1}$ $e_c^{-2}$ | Electric conversion factor |
| $k_B$ | $8.314513 \times 10^{-3}$ kJ mol$^{-1}$ K$^{-1}$ | Boltzmann constant |
| $a$ | 1.2 nm | Radius of the DNA model sphere |
| $T$ | 293 K | Temperature of the solution |
| | $4 \times 10^7$ | simulation steps for internucleosomal interaction strength 4 $k_BT$ |
| | $8 \times 10^7$ | simulation steps for internucleosomal interaction strength 8 $k_BT$ |
| | 10 nm | maximum DNA segment length |
| | 5.5 nm | nucleosome height |
| | 11 nm | nucleosome diameter |
| | 293 K | minimum temperature used for replica exchange procedure |
| | 700 K | maximum temperature used for replica exchange procedure |
| | 16 | number of temperatures used for replica exchange procedure for internucleosomal interaction strength 4 $k_BT$ |
| | 32 | number of temperatures used for replica exchange procedure for internucleosomal interaction strength 8 $k_BT$ |
| | 4 $k_BT$ and 8 $k_BT$ (inactive) | $\varepsilon$ for $E_{internuc}$ |
| | 5.5 nm | $\sigma$ for $E_{internuc}$ |
| | 665 | $a_{DNA}^{(s)}$ |
| | 665 | $a_{NUC}^{(s)}$ |
| | 120.44 | $a_{DNA}^{(b)}$ |
| | 120.44 | $a_{NUC}^{(b)}$ |
| | 219.25 | $a_{DNA}^{(t)}$ |
| | 782.85 | $a_{NUC}^{(t)}$ |
| | 1.2 kJ mol$^{-1}$ | Lennard jones $\varepsilon$ for DNA |
| | 2.0 kJ mol$^{-1}$ | Lennard jones $\sigma$ for DNA |
| | $S000$ = 1.6957 | interaction potential nucleosome s-functions |
| | $Scc2$ = –0.7641 | |
| | $S220$ = –0.1480 | |
| | $S222$ = –0.2582 | |
| | $S224$ = 0.5112 | |
| | $E000$ = 2.7206 | |
| | $Ecc2$ = 6.0995 | |
| | $E220$ = 3.3826 | |
| | $E222$ = 7.1036 | |
| | $E224$ = 3.2870 | |

**Supplementary Table 4: Oligo probe genomic coordinates and sequences**

| Chromosome | Start (hg38) | End (hg38) | Oligo name | Sequence | Dye |
|---|---|---|---|---|---|
| chr11 | 119084693 | 119084733 | ActiveA_947 | AAACTTAGCTTTGCTACAACCTTGAAATAGGCAGCATTTT | ATTO647N |
| chr11 | 119084735 | 119084775 | ActiveA_948 | AGCAAGTAGTGCAGCTTCTATGGCGCTTCCTTTGCTCTGT | ATTO647N |
| chr11 | 119084777 | 119084817 | ActiveA_949 | GGACCTCCCCATTCGACCACCCCATTCCCCAGCTGTGACA | ATTO647N |
| chr11 | 119084819 | 119084859 | ActiveA_950 | GAGGTCCCTCCCTCTGGGCGGGAATTGGAACATTGCGACA | ATTO647N |
| chr11 | 119084861 | 119084901 | ActiveA_951 | CCGCAGAGCCTCGCGTCACTTCCGGCGCCCTCCCTCGAAG | ATTO647N |
| chr11 | 119084903 | 119084943 | ActiveA_952 | CACGTGGGACCCGAGGTCGTCCTACAGTCTGACTCCTGGT | ATTO647N |
| chr11 | 119084945 | 119084985 | ActiveA_953 | GGCCCCGGGAAGCCGGGGGCTCCGGCCGGCGAGTACCGGG | ATTO647N |
| chr11 | 119084987 | 119085027 | ActiveA_954 | GCTTGGAAAGTAGGCTGTGTGTGGGTGCCGCTAAGGTCCC | ATTO647N |
| chr11 | 119085029 | 119085069 | ActiveA_955 | CACCGCCGTTGCAGCCGCATTGCCGTTACCAGACATGGCT | ATTO647N |
| chr11 | 119085071 | 119085111 | ActiveA_956 | TCGTCCAGAAGCCCAAAGTGTGCTGGTCACCGGCTCAGCA | ATTO647N |
| chr11 | 119085113 | 119085153 | ActiveA_957 | AGACGACTGAGGATGGCAACCTGGGGCCAATCGCTGCACG | ATTO647N |
| chr11 | 119085197 | 119085237 | ActiveA_959 | AACCGAGCTACTGGCCCTTTAAAAGCTACTCGGACCAAAA | ATTO647N |
| chr11 | 119085239 | 119085279 | ActiveA_960 | CAGTATCACCGCTCTCTGATTTCCCCATTCCTTCCGGAGG | ATTO647N |
| chr11 | 119085281 | 119085321 | ActiveA_961 | GGGACCGGAGTTCCGTTCCAAACAATCCTTCCACTCTTAA | ATTO647N |
| chr11 | 119085323 | 119085363 | ActiveA_962 | GTGCATCCCAGGCCTGATGGGAATCCCACCCAGATGCCCG | ATTO647N |
| chr11 | 119085365 | 119085405 | ActiveA_963 | TGTAGACGAACGTTCTTGGTCTGGGTCACTAAATCTAGAG | ATTO647N |
| chr11 | 119085407 | 119085447 | ActiveA_964 | GCATCCGCCTCAGCCCAGCCTCGAATGAAAGGACCCCGTC | ATTO647N |
| chr11 | 119085449 | 119085489 | ActiveA_965 | TGAATCAAAAGTGGAACGTGTCTTCCCAAAGGGGCCGTAT | ATTO647N |
| chr11 | 119085491 | 119085531 | ActiveA_966 | CGTTTGTGTGCAGTTCGGAGGCTTGGCTGATACTCTCTCC | ATTO647N |
| chr11 | 119085533 | 119085573 | ActiveA_967 | TGGGAGACCACTATAACACAAAAAGAAGGCGCACTTCTAA | ATTO647N |
| chr11 | 119085575 | 119085615 | ActiveA_968 | GTAGGTGTGTCACAGCTGGGGACTTGGAGCGTTGGCTGTG | ATTO647N |
| chr11 | 119085617 | 119085657 | ActiveA_969 | TGGAAAGGCCCACGGCGGCCTCCCACCCACGGTAGTAATT | ATTO647N |
| chr11 | 119085659 | 119085699 | ActiveA_970 | TTGTCTGTGCACAAGCCCAGGGCTCGGCAAGCAGGCTCGT | ATTO647N |
| chr11 | 119085701 | 119085741 | ActiveA_971 | TTCTTATCTCGGCCTGGCAGTGGCCTCCACCAGCTCTGCA | ATTO647N |
| chr11 | 119085743 | 119085783 | ActiveA_972 | CCAACAGGCTGGACAGGGGGAGATTAGCAGCTCCCCATCT | ATTO647N |

| | | | | | |
|---|---|---|---|---|---|
| chr11 | 119085785 | 119085825 | ActiveA_973 | AGGGCTCTGGAATGACTGCAAGAGCAAAGATCCCAGCCCT | ATTO647N |
| chr11 | 119085827 | 119085867 | ActiveA_974 | AGAGAGTTGGGGCCACAATTCAGATCTTCCTACTCCTAGT | ATTO647N |
| chr11 | 119085911 | 119085951 | ActiveA_976 | CACCTTTGGGACCATGTTACAGAAGCAACGGGTGGCATCA | ATTO647N |
| chr11 | 119085953 | 119085993 | ActiveA_977 | ATTGTTGTCGTCTCAAAAATCCTATCAGGTGGACAAGACA | ATTO647N |
| chr11 | 119085995 | 119086035 | ActiveA_978 | AAAACACTGAGCACTTCTATGTTGAAGCTATTGCTTTTGC | ATTO647N |
| chr11 | 119089649 | 119089689 | ActiveB_1065 | GTCCACTCTATAGAGATGGGAGAGGGGGGCAAAGAGAAAG | ATTO594 |
| chr11 | 119089691 | 119089731 | ActiveB_1066 | AGGAAGCACAGTGGGCAGGTCCTTCAAGGAGTGAACAACC | ATTO594 |
| chr11 | 119089733 | 119089773 | ActiveB_1067 | GCAAGACTCTTACTTGCAGATGGCTCCGATGGTGAAGCCA | ATTO594 |
| chr11 | 119089775 | 119089815 | ActiveB_1068 | AAAGGTTCACATGATGCCTACCCCTGCCCAAGCCCCTTAC | ATTO594 |
| chr11 | 119089817 | 119089857 | ActiveB_1069 | AGATAACCCTGAAGCAGAGGGTCAGGCCCCAAAGGGAAAG | ATTO594 |
| chr11 | 119089901 | 119089941 | ActiveB_1071 | GATCACCTCTATTCTCTCTCGATTGCCAACTCACTGGTAT | ATTO594 |
| chr11 | 119089985 | 119090025 | ActiveB_1073 | GGGTGAAAGACAACAGCATCATGAGGGTTTTCCCGCCTGG | ATTO594 |
| chr11 | 119090027 | 119090067 | ActiveB_1074 | CTCTTCTCTGGCAGGGTTTCTAGGGTCTTCCCAACAAATT | ATTO594 |
| chr11 | 119090069 | 119090109 | ActiveB_1075 | CTGGGCACATCCCACCAAGCTGCCTATCCAGGCCCCACTT | ATTO594 |
| chr11 | 119090153 | 119090193 | ActiveB_1077 | CACACTGCCCAGAGAGAAGTTACAATGAGATTTTAACACT | ATTO594 |
| chr11 | 119090195 | 119090235 | ActiveB_1078 | TCTCTGCAGCTGGGCTGCTCTTCGCAGGGAGCTGGTTCCC | ATTO594 |
| chr11 | 119090237 | 119090277 | ActiveB_1079 | CTAAAAGGATACAATACTCCTGAACTCCAGATGCGGGAAC | ATTO594 |
| chr11 | 119090279 | 119090319 | ActiveB_1080 | GCTGCTGTCTCCGTCACTCTTCCAAAAGGATCCGTCACTC | ATTO594 |
| chr11 | 119090321 | 119090361 | ActiveB_1081 | CTCCGGACTCCCAGAGCCCTCTAGACCTTGTCTTTTTCCT | ATTO594 |
| chr11 | 119090363 | 119090403 | ActiveB_1082 | GCCACTGACCCCACGGGCTGCTTGCTGGAAGCCCCTTCCA | ATTO594 |
| chr11 | 119090405 | 119090445 | ActiveB_1083 | CTGTAAATGAGTGGACGGATGAGTGCATGGAAAGACAGAC | ATTO594 |
| chr11 | 119090447 | 119090487 | ActiveB_1084 | CTTGCACTCTGAACACTTGTCTGGGGCTAAGAAAACATTA | ATTO594 |
| chr11 | 119090489 | 119090529 | ActiveB_1085 | ATGCAGGGAAATGTTTATCTTGCTCACCATTATCCCCAAT | ATTO594 |
| chr11 | 119090531 | 119090571 | ActiveB_1086 | AGTATAACTGCATTATCCCTAAGTAAGACGTAAACTCTAC | ATTO594 |
| chr11 | 119090573 | 119090613 | ActiveB_1087 | GAAGTACTCCCTATCCCTCCAGAAGTAGTCACTATTCAGT | ATTO594 |
| chr11 | 119090909 | 119090949 | ActiveB_1095 | CAAGACAATATAGTAGCATATAAAAATGGAAGTACTGACC | ATTO594 |
| chr11 | 119090951 | 119090991 | ActiveB_1096 | AATTTAGTTTTATGAAATGTACTACATATTCATTGTAAAA | ATTO594 |

| | | | | | |
|---|---|---|---|---|---|
| chr11 | 119090993 | 119091033 | ActiveB_1097 | CCCAACTTCACTTATTGGAGCACTTAGCACATACTATTTT | ATTO594 |
| chr11 | 119091035 | 119091075 | ActiveB_1098 | CTTCTGTGTTTAGATGTTCCAACTAGAAGTTTCCAGAAAA | ATTO594 |
| chr11 | 119091203 | 119091243 | ActiveB_1102 | CTGCTCTAAAAGTACTTTCCATATGTCAGCTCATTTCATC | ATTO594 |
| chr11 | 119091287 | 119091327 | ActiveB_1104 | AGCTGGCTGACTTTCAGTCATCTTCCAGCTCTCAGCTTAA | ATTO594 |
| chr11 | 119091329 | 119091369 | ActiveB_1105 | CCTCGTGCGGTTCCCTCTGCCTGAGTCTGTCTTTCCCTCT | ATTO594 |
| chr11 | 119091371 | 119091411 | ActiveB_1106 | TGTGGAGAGTTGCTAAGGACCACAGTGTCGGGCAATCTGG | ATTO594 |
| chr11 | 119091413 | 119091453 | ActiveB_1107 | TGCTCGTCCAGCTTCCGAAGCCGGGTGTTGAGGTTTCCCC | ATTO594 |
| chr11 | 119091455 | 119091495 | ActiveB_1108 | TGCAGGCCAGCTGTTGCCAGGATGATGGCACTGAACTCCT | ATTO594 |
| chr11 | 119091497 | 119091537 | ActiveB_1109 | GCAGGCCCTACCTGCCCCACCCGGTTGTGCCAGCCCATGC | ATTO594 |
| chr11 | 119094857 | 119094897 | ActiveC_1189 | ACCACCGCCCTCATGGAAAGAGCTGAGCCGCTTCAGACTG | ATTO647N |
| chr11 | 119094899 | 119094939 | ActiveC_1190 | GCCAGGCCTCCCCATGCCACCACAAAGGCCCTTTTAAGGG | ATTO647N |
| chr11 | 119094941 | 119094981 | ActiveC_1191 | GGCCTCCCAGGAGTACTAAGAGGGCCCGCGCCGCGGCCGG | ATTO647N |
| chr11 | 119094983 | 119095023 | ActiveC_1192 | CGTGGGGCCGAAGGCGCCCTCGGGCGGCAAGAAGGCCACC | ATTO647N |
| chr11 | 119095067 | 119095107 | ActiveC_1194 | GCTGCTGGGCGGCGTGACGATCGCCCAGGGAGGCGTCCTG | ATTO647N |
| chr11 | 119095277 | 119095317 | ActiveC_1199 | CCCAGTGGGCCGTGTACACCGGCTGCTGCGGAAGGGCCAC | ATTO647N |
| chr11 | 119095361 | 119095401 | ActiveC_1201 | CGCTAGCATGTCGGGCCGCGGCAAGACTGGCGGCAAGGCC | ATTO647N |
| chr11 | 119095403 | 119095443 | ActiveC_1202 | TCTGTTCTAGTGTTTGAGCCGTCGTGCTTCACCGGTCTAC | ATTO647N |
| chr11 | 119095445 | 119095485 | ActiveC_1203 | CCTCGCGGCGCGCGCGCGACAGCAGTTACACTGCGGCGGG | ATTO647N |
| chr11 | 119095487 | 119095527 | ActiveC_1204 | CAGGAGGGCGCAGAGGTGTGTCCTGGGGGCTTATAAAGGC | ATTO647N |
| chr11 | 119095529 | 119095569 | ActiveC_1205 | ACCGCAACCAACCGGAGGCGGGTATTGGAGAAAAGAGCCA | ATTO647N |
| chr11 | 119095571 | 119095611 | ActiveC_1206 | GCGGGCGCGCTCCCTCAGCCGAGCCGCGACACCCAAGGGT | ATTO647N |
| chr11 | 119095613 | 119095653 | ActiveC_1207 | CCTTTTCTCCCCGGGCAAAAGGTTTTCAAATTCGACCAAT | ATTO647N |
| chr11 | 119095655 | 119095695 | ActiveC_1208 | TCAGCGGATAGTTGGCAGTCTGCGTGCTACCCCTCCTCCT | ATTO647N |
| chr11 | 119095697 | 119095737 | ActiveC_1209 | TGTAGTGTTTAAATCTCGCGCGCTTTACAAGGATTGGCTA | ATTO647N |
| chr11 | 119095739 | 119095779 | ActiveC_1210 | CTCAGACTCCACCCCTTTGGTCTTCCGCTTCTGGTTTCCG | ATTO647N |
| chr11 | 119095781 | 119095821 | ActiveC_1211 | ATCCTACGCCGGCACAGATTTCCCAGACGCTCTCTAGGTG | ATTO647N |
| chr11 | 119095823 | 119095863 | ActiveC_1212 | GAGGGCTGCAGTAGCTATAATATTGACCCCTTTCCTTTAA | ATTO647N |

| | | | | | |
|---|---|---|---|---|---|
| chr11 | 119095865 | 119095905 | ActiveC_1213 | TCAGGTAGGAGAAGGGAGTAGGGAGTGACTGGACCTGGAA | ATTO647N |
| chr11 | 119096285 | 119096325 | ActiveC_1223 | GCATGGAGGGCCACAAAATTGAAGTTATCCTGTTTCCTGG | ATTO647N |
| chr11 | 119096369 | 119096409 | ActiveC_1225 | CAGACCCCGTCTAACAAGAAGTGAACAGAACTGAGACCAA | ATTO647N |
| chr11 | 119096411 | 119096451 | ActiveC_1226 | GGGGGTACTTCCTCCAACAGGATGCCTATCTCTCTGCCCC | ATTO647N |
| chr11 | 119096453 | 119096493 | ActiveC_1227 | ATTTCCTATATGGGCATTTTTGGGTAGATTGGGAGGGGTG | ATTO647N |
| chr11 | 119096537 | 119096577 | ActiveC_1229 | GATTAGTAAGAACTAAGCAGGGGGCCACATGCTCTCAATG | ATTO647N |
| chr11 | 119096579 | 119096619 | ActiveC_1230 | GCTCCACGGTGACATGTCATTTGATTGTAAATTAAGTGTT | ATTO647N |
| chr11 | 119096621 | 119096661 | ActiveC_1231 | CTGGTTTTTGGGATAGAACTTGGGCCAGGGCTAGGAACAC | ATTO647N |
| chr11 | 119096663 | 119096703 | ActiveC_1232 | TATCCACACTCACATTTCAGAGTCCTGACTCTCAAGGAAC | ATTO647N |
| chr11 | 119096705 | 119096745 | ActiveC_1233 | TTGCAGCCTCATAGGGTGGGATACAGCAGCTTTTTTTGCA | ATTO647N |
| chr11 | 119096789 | 119096829 | ActiveC_1235 | CGTTTCTTGCCCTCTGCTGACTACTGATTGGATTTTACCT | ATTO647N |
| chr11 | 119096873 | 119096913 | ActiveC_1237 | TTGCCCAGGCCTCTCTCACTCTTCATACTCCTCCAGATTT | ATTO647N |
| chr11 | 119100569 | 119100609 | ActiveD_1325 | TGCCCCTATGGCACCTACTTCAGGGAACCCTTCCTGGTGC | ATTO594 |
| chr11 | 119100611 | 119100651 | ActiveD_1326 | CTTGGCCTGCATCTGGACTTGGGTAGGTAGTCCTACCACT | ATTO594 |
| chr11 | 119100653 | 119100693 | ActiveD_1327 | GGCAACACGACCATTGTGGTGCCCAAGCCCTTCCGCCCGA | ATTO594 |
| chr11 | 119100695 | 119100735 | ActiveD_1328 | GCTGCCTCACTACCTCTCCTCATGGTCTATTTCACCAACT | ATTO594 |
| chr11 | 119100737 | 119100777 | ActiveD_1329 | GTACTGAATCTGCGCTGGCGCCATAAGCTGCTGCTACCTA | ATTO594 |
| chr11 | 119100779 | 119100819 | ActiveD_1330 | CTTGCCATCTGCTGCATGATCTTCCTGGGCTTTGCGGATG | ATTO594 |
| chr11 | 119100821 | 119100861 | ActiveD_1331 | TGCCCGAGCCTCCCCCAGTTTGTGGCCCTGATAGGTGCCC | ATTO594 |
| chr11 | 119100863 | 119100903 | ActiveD_1332 | ATTCTTAAAAGGTGGAATGGGAGCAGGCTTGAGTCATGGA | ATTO594 |
| chr11 | 119100905 | 119100945 | ActiveD_1333 | GGCTTGTGGGGAGGGGCTAAGAAATTATCAGAAAAGACAG | ATTO594 |
| chr11 | 119100947 | 119100987 | ActiveD_1334 | GGCTGGGACCTGGGAGGTACCTGAGAGAACTGGGGTTATT | ATTO594 |
| chr11 | 119100989 | 119101029 | ActiveD_1335 | CCCCACCATGAAGTAAGTGGGTTCGTGGGGGTGATTGCCT | ATTO594 |
| chr11 | 119101031 | 119101071 | ActiveD_1336 | CCCTTCCTGAACTGCTTTGTGAAGGAGCAGTGTAAGGCAT | ATTO594 |
| chr11 | 119101073 | 119101113 | ActiveD_1337 | GGTGCTGTTTTCCTTATCATCCTCTTCTGCTTCATCCCTT | ATTO594 |
| chr11 | 119101115 | 119101155 | ActiveD_1338 | TTGCTGCTGGGCCACAGCCCAGAATCCCAGGGAGTGATCA | ATTO594 |
| chr11 | 119101157 | 119101197 | ActiveD_1339 | TAGTGAGTGACCACGCCCCCTTTCCTCTTCCCCCTCGCCC | ATTO594 |

| | | | | | |
|---|---|---|---|---|---|
| chr11 | 119101199 | 119101239 | ActiveD_1340 | CGGGAATGAGAAGACCACTTTGGGTTACTGTAACACCTGC | ATTO594 |
| chr11 | 119101241 | 119101281 | ActiveD_1341 | TTCCTCCCTCCGCCCCCCTCACCACTTACCAGAATAAAAA | ATTO594 |
| chr11 | 119101283 | 119101323 | ActiveD_1342 | GTAGGGACCCTTGGGTATATCTGGGACTCTGGCAGTGGTG | ATTO594 |
| chr11 | 119101325 | 119101365 | ActiveD_1343 | AGAGGTGATGAGCAGAACTTACTCGCATTGGGGAAAGGAT | ATTO594 |
| chr11 | 119101367 | 119101407 | ActiveD_1344 | CACGTGAAGACTCAGAACTAACCCAGGCAGCCTGGAACTC | ATTO594 |
| chr11 | 119101409 | 119101449 | ActiveD_1345 | AGGAGTGGCTAGGGCAGGGGCGGGAACCGGGGTGCTTGAC | ATTO594 |
| chr11 | 119101451 | 119101491 | ActiveD_1346 | AGCAGCGGCACACGGGTCCGGGCAGGGGGCAAGGGCTAAG | ATTO594 |
| chr11 | 119101493 | 119101533 | ActiveD_1347 | CTGTGGTCAGGACCTCAACAAAACCAGCCGACAGCAGATG | ATTO594 |
| chr11 | 119101535 | 119101575 | ActiveD_1348 | CCTCATCCCGGCCTTCCGGGGCCACTTCATTGCTGCGCGC | ATTO594 |
| chr11 | 119101577 | 119101617 | ActiveD_1349 | CAATTTGATCGTCTCGCTGCTGGGATTTGTGGCCACAGTC | ATTO594 |
| chr11 | 119101619 | 119101659 | ActiveD_1350 | CACCATGTGGGCCTTCTCGGAATTGCCCATGCCGCTGCTG | ATTO594 |
| chr11 | 119101661 | 119101701 | ActiveD_1351 | CCCGTTACCTGAAGAGGCGGCGGAGCCGGGCCCCTGACCG | ATTO594 |
| chr11 | 119101703 | 119101743 | ActiveD_1352 | CCACTCCATACTGCTGAGGCCTCAGGACTGCTGCTCAGCT | ATTO594 |
| chr11 | 119101745 | 119101785 | ActiveD_1353 | TTGCTCAGATTGGTGTGGGAAGAGCCTGCCTGTGGGGAGC | ATTO594 |
| chr11 | 119101787 | 119101827 | ActiveD_1354 | CCCCGGTGTCCCGACGGCGGCTCAAGTCAGAGTTGCTGGG | ATTO594 |
| chr11 | 119105819 | 119105859 | ActiveE_1450 | AAGACCCTCTTCCCCCAAGAATCCTTCAGGAACCCTCAGA | ATTO647N |
| chr11 | 119105861 | 119105901 | ActiveE_1451 | CAGGTTATGCAGGAAGTTGTCCAGGTAGAAGCTTTCCCTA | ATTO647N |
| chr11 | 119105903 | 119105943 | ActiveE_1452 | GAGGAAGCTGTCAGTGGGAAGAGCAGCTAGGGTTTAGGGG | ATTO647N |
| chr11 | 119105945 | 119105985 | ActiveE_1453 | ATCAAGGGAGGAGAGAACTCAGGGCCAGAACCAGGGATGC | ATTO647N |
| chr11 | 119105987 | 119106027 | ActiveE_1454 | AGAGTCTGGGTTTCTAATCTCAAAGGGAGGAGAAGACATG | ATTO647N |
| chr11 | 119106029 | 119106069 | ActiveE_1455 | CCCATGCATTGTCACTGGACTGCCAAATCAGTGGAGATTC | ATTO647N |
| chr11 | 119106071 | 119106111 | ActiveE_1456 | GGGGAGAGGTGGGGGTGAATGAGGTAACCTCCTGTGGCCC | ATTO647N |
| chr11 | 119106113 | 119106153 | ActiveE_1457 | GGGGTAGCCTGGGCCTCCCTAGATTTCCCTGTAAGGAATG | ATTO647N |
| chr11 | 119106155 | 119106195 | ActiveE_1458 | TGACAGGTCAGCAACAGGTTTGTCTGAGCCAGAGGAGGTG | ATTO647N |
| chr11 | 119106197 | 119106237 | ActiveE_1459 | CAATGCCTTCTGAAGTGTGGGATGGCAGGGCTAGACAGAA | ATTO647N |
| chr11 | 119106239 | 119106279 | ActiveE_1460 | GTAGTTGTCTGAGGAATGAGCACATGAGAATAAGAGAATA | ATTO647N |
| chr11 | 119106281 | 119106321 | ActiveE_1461 | GGTAGGATGGAAAGGTGTTAAGTATCTTTCTCCCAAACCC | ATTO647N |

| | | | | | |
|---|---|---|---|---|---|
| chr11 | 119106323 | 119106363 | ActiveE_1462 | ACATGTGGGAAGGTCTTTGGCACACAGTCCCTTTCTTAAA | ATTO647N |
| chr11 | 119106365 | 119106405 | ActiveE_1463 | ATCCTGGGGGAATTCACATTCTCCCAAGAGCCCCTTCTTC | ATTO647N |
| chr11 | 119106407 | 119106447 | ActiveE_1464 | TGAGGACAGGGGAAGGAAATACCTGCAAAAGGATTCAAAG | ATTO647N |
| chr11 | 119106449 | 119106489 | ActiveE_1465 | GTTGATTCAACCTCTACAGGACTGGGGCTCATCAATGCTT | ATTO647N |
| chr11 | 119106491 | 119106531 | ActiveE_1466 | CAACCTCTGGCCTCAGAAGCCATCTTTTCCTAGACTCAAG | ATTO647N |
| chr11 | 119106533 | 119106573 | ActiveE_1467 | TCCCAACAAGCTCCCAAGAAATGGTGCTGTAGGTACACTG | ATTO647N |
| chr11 | 119106575 | 119106615 | ActiveE_1468 | ATAGATTCAGCCACAAGACCTTGCCACTCTAAGTATGTGA | ATTO647N |
| chr11 | 119106617 | 119106657 | ActiveE_1469 | TGGTAATGACAAGGGAGCTGGCAATGTAGCACTGGACTAT | ATTO647N |
| chr11 | 119106659 | 119106699 | ActiveE_1470 | CCATAGTAACAGCTCCTGCCTATGGAGCACGTGTAGAGGG | ATTO647N |
| chr11 | 119106743 | 119106783 | ActiveE_1472 | CCCATTAACCTCCTTGCTCGTGTAGGTTTGTTCTGGAGAT | ATTO647N |
| chr11 | 119106785 | 119106825 | ActiveE_1473 | TCATCACCCTCCTTCCTCTTGGAGCTTGGACCCAGAAACA | ATTO647N |
| chr11 | 119106827 | 119106867 | ActiveE_1474 | AGAGGTTGAGAGTAGAACACCGAATTTGAATTCCCCCCAC | ATTO647N |
| chr11 | 119106911 | 119106951 | ActiveE_1476 | GACGCAGAGCGTCCCTCCCAGATAGAAGTGTCTGGCAGAG | ATTO647N |
| chr11 | 119106953 | 119106993 | ActiveE_1477 | TAGCTTCCCATTTCCTGGGGGAAGCTAGGCCCAGGGTAAG | ATTO647N |
| chr11 | 119106995 | 119107035 | ActiveE_1478 | TCACCGCCCCCTTCTCCCAACACCCCCAGACTCAGCGACT | ATTO647N |
| chr11 | 119107037 | 119107077 | ActiveE_1479 | CGGACGCCTGTGAGGTTAAGAAGGAGGTGACCCCTGTGGA | ATTO647N |
| chr11 | 119107079 | 119107119 | ActiveE_1480 | GCTGCCTTTGCTTTCCCCTATGCTTGTTAGATTTCCACAG | ATTO647N |
| chr11 | 119107121 | 119107161 | ActiveE_1481 | CACACTCAGCTCAAGGCAAGAGTAACTTCACTTACCTAAG | ATTO647N |
| chr11 | 55812948 | 55812988 | InactiveA_64 | AAAGTTCCAATGACACTGGAAACTAAAACAACAGCAAGAA | ATTO647N |
| chr11 | 55812990 | 55813030 | InactiveA_65 | TGAAATTATTATGTTTTTCATCTATCATATAACATATTAT | ATTO647N |
| chr11 | 55813284 | 55813324 | InactiveA_72 | ACAGAGATTAAAAGTCCTCTTATAAGTGTCAGTAAGCATG | ATTO647N |
| chr11 | 55813368 | 55813408 | InactiveA_74 | AATTCAGAAATGTGGTTGGGGTACATATGGAAAGTGAAAT | ATTO647N |
| chr11 | 55813410 | 55813450 | InactiveA_75 | AGTCACAAAGAAAGCATCAATGGTGCATATAATTCAATAT | ATTO647N |
| chr11 | 55813494 | 55813534 | InactiveA_77 | TTCACACGGTATCATAGTATAAAATTTTTCTGAATCTTGT | ATTO647N |
| chr11 | 55813578 | 55813618 | InactiveA_79 | CACTTAAATAAGTACTTGTCTTTATTAAGTTTGCTTCCAC | ATTO647N |
| chr11 | 55813620 | 55813660 | InactiveA_80 | TGTGATTCTTTTCTCATTTGCTGGCAAACTTTGTTTTAAT | ATTO647N |
| chr11 | 55813746 | 55813786 | InactiveA_83 | AAGAATGTTCACAGTGACTCTGTTATGGCTTTTAGTGGTA | ATTO647N |

| | | | | | |
|---|---|---|---|---|---|
| chr11 | 55813872 | 55813912 | InactiveA_86 | GTTTAGAGATAAAAGGTTTCCTTTTAGAGAAAATTTGGAA | ATTO647N |
| chr11 | 55813998 | 55814038 | InactiveA_89 | AGAGACAATCTGATTAACTGTTGACAGTATTCTAGCTTCC | ATTO647N |
| chr11 | 55814040 | 55814080 | InactiveA_90 | TAAACCAGTGGCAGCAATTTCAAATCTCTTCCTATGTGTC | ATTO647N |
| chr11 | 55814124 | 55814164 | InactiveA_92 | TTTTTAGTATCTAGATAGAAACATACATGAGGCCTTTAAA | ATTO647N |
| chr11 | 55814166 | 55814206 | InactiveA_93 | GACAACCATTAGTAGTAATCAATGATTTCTTATATTTTTT | ATTO647N |
| chr11 | 55814376 | 55814416 | InactiveA_98 | CCTAATGATTGTTATGTTCATTGAAGTCACCTTGATCCAC | ATTO647N |
| chr11 | 55814502 | 55814542 | InactiveA_101 | GAGAAGCATATGATAACTGGTTGATGACAACTAGTTGTCC | ATTO647N |
| chr11 | 55814544 | 55814584 | InactiveA_102 | TTCTTCATTGATTTTTCTGACTACTGTTGGTTGCAGTCCA | ATTO647N |
| chr11 | 55814586 | 55814626 | InactiveA_103 | TATTAGCTCAGGTTCATTCTTTTAGTGTCTTTTAGGCAAA | ATTO647N |
| chr11 | 55814670 | 55814710 | InactiveA_105 | TGTATTTGGCTTAGTGCTTCATTTTGTGTCATTGTATATA | ATTO647N |
| chr11 | 55814796 | 55814836 | InactiveA_108 | ATTCAGATAAATATGGTTGTTATAAAAGTTTACTTTGTTA | ATTO647N |
| chr11 | 55814838 | 55814878 | InactiveA_109 | ATTGTCTCAAAGGCATATTTTTTCCAGTTAGTTTGAATCA | ATTO647N |
| chr11 | 55814922 | 55814962 | InactiveA_111 | CACTCATGCCAAAACTGCACCTAACAAAATCATCATTATA | ATTO647N |
| chr11 | 55814964 | 55815004 | InactiveA_112 | TTATCACTTTTTGATTGGAGAGTCTTTAATGTTTGAACCA | ATTO647N |
| chr11 | 55815132 | 55815172 | InactiveA_116 | GCAGGCCAAACTATCGACTTCTGATTAGAGTATTTTCATT | ATTO647N |
| chr11 | 55815174 | 55815214 | InactiveA_117 | TCCCTTCAGTTCCTCCAGGGCATGGCGATGCTGAAGTACA | ATTO647N |
| chr11 | 55815216 | 55815256 | InactiveA_118 | GTCTTCCCGCTCCACCCAGTTAGATCCCACTTGATTCCCT | ATTO647N |
| chr11 | 55815258 | 55815298 | InactiveA_119 | AAGCAGACCTAGAGAAAAAGTTGAAATTTGAGCGCATTGT | ATTO647N |
| chr11 | 55815342 | 55815382 | InactiveA_121 | TATGGGAAGATCAGATAAAATTAAGAACTCATGTAAGTAT | ATTO647N |
| chr11 | 55815384 | 55815424 | InactiveA_122 | AATTAGAAAGCACTCTTTCTAAACAATTGGCCTATTGGCA | ATTO647N |
| chr11 | 55815426 | 55815466 | InactiveA_123 | TGAGCCACCATGCTCCGCCCAAGCCTTCCCTATTAAAAAA | ATTO647N |
| chr11 | 55818240 | 55818280 | InactiveB_190 | CAAGAAAGATGTTAATTTACATGCAAGACCAACATATTCA | ATTO594 |
| chr11 | 55818282 | 55818322 | InactiveB_191 | ATATCTATTCACATAACCCACCCAAATTTTTAGACGGCCT | ATTO594 |
| chr11 | 55818324 | 55818364 | InactiveB_192 | GCTATCATATCTGTAATCTCTTTCTCTATCATAATAGACA | ATTO594 |
| chr11 | 55818450 | 55818490 | InactiveB_195 | TATGAAGGAGAAGAACTAAAAACTTGAGATAGTACAGCAA | ATTO594 |
| chr11 | 55818492 | 55818532 | InactiveB_196 | CCTTCAAGACACATTATATATTCATTGTGTCCATCTATTA | ATTO594 |
| chr11 | 55818576 | 55818616 | InactiveB_198 | GTGTAAGAGTCTGACTTGCAAACTTCCCTTTTCTTCTAGG | ATTO594 |

| | | | | | |
|---|---|---|---|---|---|
| chr11 | 55818702 | 55818742 | InactiveB_201 | TAAGCAGACTAGAGAATCTACATAGTCCAGTGACAGTTAC | ATTO594 |
| chr11 | 55818744 | 55818784 | InactiveB_202 | TACTAACACATTCCCTTCATTAATAACCATACTTTCAATT | ATTO594 |
| chr11 | 55818786 | 55818826 | InactiveB_203 | CCTTTACCCGCACCAAACTCTGCTCCATTATCTAGGTCGA | ATTO594 |
| chr11 | 55818828 | 55818868 | InactiveB_204 | CTGTGACTAGGGTTTTGATCCTTGAACTCCTAGACATGTA | ATTO594 |
| chr11 | 55818870 | 55818910 | InactiveB_205 | TGAATTTCCTATCGATATGCCCAGGAAGGCCAATCTACAC | ATTO594 |
| chr11 | 55818912 | 55818952 | InactiveB_206 | TTCTTTATTCTATAGGTGACAAAGTAGAAGCCAGAGAAGC | ATTO594 |
| chr11 | 55818954 | 55818994 | InactiveB_207 | AAATAATAAATATCGAATAGTCAATACAGATACCCTTGTT | ATTO594 |
| chr11 | 55818996 | 55819036 | InactiveB_208 | AATCAGGCTGTACTAATTATCAATCCTTCCTCATTGCTTG | ATTO594 |
| chr11 | 55819038 | 55819078 | InactiveB_209 | CAAAATACAGCTGAACTAATTGTTTCAGAGCGTATGCTCC | ATTO594 |
| chr11 | 55819080 | 55819120 | InactiveB_210 | ATATAGCAGTTTAATGTTTGTCTGACCCAGAAAATTTGAC | ATTO594 |
| chr11 | 55819164 | 55819204 | InactiveB_212 | TAGTTTTCACATAATTTGTTTATACGATGACATTAACTAT | ATTO594 |
| chr11 | 55819248 | 55819288 | InactiveB_214 | CTACATAAATGATCTTCCTCGTATAATATGATGACTGATT | ATTO594 |
| chr11 | 55819500 | 55819540 | InactiveB_220 | ACGGCTCCTGAAGGATGAAGCAAATATCTCTTCCGGAATC | ATTO594 |
| chr11 | 55819584 | 55819624 | InactiveB_222 | CTGTTTCTTCGAATCTGAAAGTTAGTTCTGAAAACAAAAA | ATTO594 |
| chr11 | 55819626 | 55819666 | InactiveB_223 | ACGTGGTCCCACTTGTATTTCTATCAGTCAGCAGCATCGC | ATTO594 |
| chr11 | 55819710 | 55819750 | InactiveB_225 | GCACAGTGACATTGTAGATGGCCAGAAAAACCAGGAAGAG | ATTO594 |
| chr11 | 55819752 | 55819792 | InactiveB_226 | TGGGGTTGATTTTGATGATCACAATCAACCCAATATTCCC | ATTO594 |
| chr11 | 55819878 | 55819918 | InactiveB_229 | CTAAAAATGAAATGGTTCTGTCTTTGACAACAAGGTTCAC | ATTO594 |
| chr11 | 55820004 | 55820044 | InactiveB_232 | GGGACATGTTAACTGTGTAGAGCAGAGGGTTGCAAATGGC | ATTO594 |
| chr11 | 55820046 | 55820086 | InactiveB_233 | CCCAGGCATAGGATCCCACAACCAGCAGCACGCAGAGTTT | ATTO594 |
| chr11 | 55820172 | 55820212 | InactiveB_236 | TGTAAGTATCAGAGCAAGAAAGGGAGAGTAGTGAGGAGAA | ATTO594 |
| chr11 | 55820214 | 55820254 | InactiveB_237 | TTTCATTAAAGGTGGCAAGAAAGAATAGCAGCCACTGGTT | ATTO594 |
| chr11 | 55820256 | 55820296 | InactiveB_238 | CAATGAACGCATAAGATGTGAGAACGATGAGTAGTGTGCT | ATTO594 |
| chr11 | 55820298 | 55820338 | InactiveB_239 | TGCGGCGCCCACTGACTGAACGCATCTTGAGGATGGTTAC | ATTO594 |
| chr11 | 55823448 | 55823488 | InactiveC_314 | CGGGGGTTACAGACCACCTTGGGCATAATGGAAGAACCTT | ATTO647N |
| chr11 | 55823532 | 55823572 | InactiveC_316 | ATAGAACCTTGCCCAGGTGTGGTGATTAATGCCCCTGATC | ATTO647N |
| chr11 | 55823574 | 55823614 | InactiveC_317 | CTTTTCATTGAAAAGAATCGCAGTTGACATGTACTATAGC | ATTO647N |

| | | | | | |
|---|---|---|---|---|---|
| chr11 | 55823616 | 55823656 | InactiveC_318 | CAAATCCATGATATGACTTGTTTTTCTATGGTGTTTGTAA | ATTO647N |
| chr11 | 55823658 | 55823698 | InactiveC_319 | TGTTATTCATGAGGGTTGAAGTTGTCTTTAATCGAAACAC | ATTO647N |
| chr11 | 55823742 | 55823782 | InactiveC_321 | GATTTCTCTAATCCATTCATACTTACCATGTGAATCACTC | ATTO647N |
| chr11 | 55823784 | 55823824 | InactiveC_322 | ATTCTCACAACAGCTTTTCTCTCGCTCTTTAAAAAAGCAT | ATTO647N |
| chr11 | 55823826 | 55823866 | InactiveC_323 | CTGCATTGTAGGAGGCTTTCTGGGTTTCCTTTTGTAACTA | ATTO647N |
| chr11 | 55823910 | 55823950 | InactiveC_325 | AAGAGAAACCAAACATTTCAATAATTCCATTGATATAAGC | ATTO647N |
| chr11 | 55823952 | 55823992 | InactiveC_326 | CAACCTGAATGAGCAGTGTTCAAAGCACAGTCTGTGTGCC | ATTO647N |
| chr11 | 55823994 | 55824034 | InactiveC_327 | TCTCCCCAAGTTCTGCTTCCTTCTTGACTAATACAGCAAT | ATTO647N |
| chr11 | 55824036 | 55824076 | InactiveC_328 | CTATTCAATGCTAGTGGCCATTAAATGGGAATATCTCAAG | ATTO647N |
| chr11 | 55824078 | 55824118 | InactiveC_329 | GGAAGAGCAGAGCTCCAGAGTGGTACCGGGGGTGCAGACT | ATTO647N |
| chr11 | 55824120 | 55824160 | InactiveC_330 | TCAAGTGTGGCTTTTTGGCTTGTGCCATCTGTGCAACCAT | ATTO647N |
| chr11 | 55824162 | 55824202 | InactiveC_331 | AACTCACCCACAGTCACACATAGAATCCCTGATAGAGACA | ATTO647N |
| chr11 | 55824204 | 55824244 | InactiveC_332 | ATTTCCCCTTTCAAAATAAGGTTATAGAGGCACAGCAGGT | ATTO647N |
| chr11 | 55824246 | 55824286 | InactiveC_333 | ACACATTTTATCTTCATAAAACTGGTTGACATTAGTACTG | ATTO647N |
| chr11 | 55824288 | 55824328 | InactiveC_334 | GTACTGCTCATGTACCAGATACTATTCTAAATACAATTAT | ATTO647N |
| chr11 | 55824330 | 55824370 | InactiveC_335 | GAATACTAATAATTGCTGATACTAATACTAATAATTTACT | ATTO647N |
| chr11 | 55824372 | 55824412 | InactiveC_336 | AAATACCAGAAAGGGGTGGGGATAAGGAGTCCCTTTTACT | ATTO647N |
| chr11 | 55824456 | 55824496 | InactiveC_338 | TAAATAGGCATTATAAAATTTGTTTCACACTCAAAAGCTC | ATTO647N |
| chr11 | 55824498 | 55824538 | InactiveC_339 | TGAGCCAGTACTCACATTGCCATTTCATGTTATGAAATTA | ATTO647N |
| chr11 | 55824540 | 55824580 | InactiveC_340 | AAATTGCATAACATTGTATGCCACATCCAAGCATTGCCTC | ATTO647N |
| chr11 | 55824582 | 55824622 | InactiveC_341 | CATATTATTATAATGTAACACAATTACATTCAGACTTCAT | ATTO647N |
| chr11 | 55824666 | 55824706 | InactiveC_343 | CAAGGAAACCATCCTATCCTGGGACTCCAACAATTATTGA | ATTO647N |
| chr11 | 55825002 | 55825042 | InactiveC_351 | AAATTATCTAATACCATTACCATCCTTTTGAATTCCCCTT | ATTO647N |
| chr11 | 55825170 | 55825210 | InactiveC_355 | CTAATAGCTGTTATAAGATGTTTGGTTTTACAGATTAAAT | ATTO647N |
| chr11 | 55825212 | 55825252 | InactiveC_356 | AAGTGCTTATGCCAATGTCTTCACACTGCTTTACATCTCA | ATTO647N |
| chr11 | 55825296 | 55825336 | InactiveC_358 | TAGGAAAATCTCCATGCAGTCTTAACCAAGTCGCATGGAC | ATTO647N |
| chr11 | 55825338 | 55825378 | InactiveC_359 | CCGTGGGCCTCCTACTGCCACCCTCAGCACTAGGATGACT | ATTO647N |

| | | | | | |
|---|---|---|---|---|---|
| chr11 | 55828698 | 55828738 | InactiveD_439 | TAGAGTCCGCTGCAATTATTATTTTCGTCATCTATCAAAT | ATTO594 |
| chr11 | 55828740 | 55828780 | InactiveD_440 | AGCTCTTTTTATGTGATTCCAGCTTCCTTAATGCAATTTC | ATTO594 |
| chr11 | 55828782 | 55828822 | InactiveD_441 | GAGAGAGAGACAACTAATAATTATGTTCCAGGGATTATTC | ATTO594 |
| chr11 | 55828950 | 55828990 | InactiveD_445 | TTTAGGGTTAGTATGAAGATAAGCAGTAAAGATAAAGTGA | ATTO594 |
| chr11 | 55828992 | 55829032 | InactiveD_446 | TAAAAGTCATAAGTGTCAGTAAGCATCTCTTCATCAATAG | ATTO594 |
| chr11 | 55829118 | 55829158 | InactiveD_449 | GCATTAATGGTGCATATAATTCAACAGGGAATTAAGAAAT | ATTO594 |
| chr11 | 55829160 | 55829200 | InactiveD_450 | AATGAGGAAATTGACATAAGTCGTAATGGAATCACAAATA | ATTO594 |
| chr11 | 55829244 | 55829284 | InactiveD_452 | TCGGTTTTGAAAAACTCAAACCTATTGCTTTTCTAATATT | ATTO594 |
| chr11 | 55829286 | 55829326 | InactiveD_453 | AATTACTTGACAAATTAAAAAACCAGGGTCAAAACACCAA | ATTO594 |
| chr11 | 55829412 | 55829452 | InactiveD_456 | CACTGAGTCTCTTCCTGATGTCAGTAAGTACACATATGCT | ATTO594 |
| chr11 | 55829454 | 55829494 | InactiveD_457 | AGTTAAATAAACCCCAAGCTTCAATGACTTCATTTAGATA | ATTO594 |
| chr11 | 55829496 | 55829536 | InactiveD_458 | TTAAAAACTTCTCATATGCCACTGATGATGAAGTAGATAA | ATTO594 |
| chr11 | 55829580 | 55829620 | InactiveD_460 | TAATCTGCATAGTAATCCCAATATGCTTACCATAAATTCA | ATTO594 |
| chr11 | 55829622 | 55829662 | InactiveD_461 | ACAATGTGGACCAACTCACAGTTCTTGAAGTCTCTCGAAA | ATTO594 |
| chr11 | 55829664 | 55829704 | InactiveD_462 | ATTTGTTTTCCTAAGTAGACCATGACTGCTAGAACATTCG | ATTO594 |
| chr11 | 55829706 | 55829746 | InactiveD_463 | GCCTCTAATCGCTAACATTTCATGAATTAGATCCTGCAGA | ATTO594 |
| chr11 | 55829748 | 55829788 | InactiveD_464 | ACAATCTTTAAATAATACAGCATCTCCTGTGTGCTCTTTT | ATTO594 |
| chr11 | 55829790 | 55829830 | InactiveD_465 | TTGCCCTCTAATATCAAGAGCAATAGTTACTTCCTAAAGT | ATTO594 |
| chr11 | 55829832 | 55829872 | InactiveD_466 | TCCACGAAGCTCTGAGGTTCTGGGGACAATGTTTCTTCCC | ATTO594 |
| chr11 | 55829874 | 55829914 | InactiveD_467 | TGTCTCTTCCTCAAGGGATTAACACCGGCGGACTGAGCTG | ATTO594 |
| chr11 | 55829916 | 55829956 | InactiveD_468 | AGTCCTAACCCAGCCTCAGAGGCCCTGACACACTAAGCGT | ATTO594 |
| chr11 | 55829958 | 55829998 | InactiveD_469 | GGACCATGTCTGAGCGTGCCACAGGACACTGGAAATCTGC | ATTO594 |
| chr11 | 55830000 | 55830040 | InactiveD_470 | CCTCTTCAGAGGCAGGAGCAGCCATCTAGCACAGCCTTCT | ATTO594 |
| chr11 | 55830042 | 55830082 | InactiveD_471 | AGCCACAGTGGATACTTGGTGTGTTCCAGCTTTCCATGAT | ATTO594 |
| chr11 | 55830168 | 55830208 | InactiveD_474 | TTTAGATCCGCCAGAGGAAGTAAGGGGGTAAGGGGAGGAG | ATTO594 |
| chr11 | 55830210 | 55830250 | InactiveD_475 | TGCAAACTGTATTCCCCAGATGTCCTATCTAGCTGCCTTC | ATTO594 |
| chr11 | 55830252 | 55830292 | InactiveD_476 | CCACTCTCCTAGTCCTTATGCTGTGATCTTGCTGCTGGGG | ATTO594 |
| chr11 | 55830294 | 55830334 | InactiveD_477 | CCTCAGAAGCTGTGTTTGTTGAACCTAGTCTCATTTTCAT | ATTO594 |

| | | | | | |
|---|---|---|---|---|---|
| chr11 | 55830336 | 55830376 | InactiveD_478 | AGAGGGTTTTCCTTGTACCCTGTGCATTGTACACCGCAAG | ATTO594 |
| chr11 | 55830378 | 55830418 | InactiveD_479 | GTTCAGATACAGAAAGATGTACACACTCTAGGTAATAGGA | ATTO594 |
| chr11 | 55833528 | 55833568 | InactiveE_554 | TTGCATTTGAATCCATACCATATTTAATCTATTCTAAGTC | ATTO647N |
| chr11 | 55833570 | 55833610 | InactiveE_555 | CATTGTTGTTCTTTTTCCCAACAGAATTCTAGCAAAGATC | ATTO647N |
| chr11 | 55833654 | 55833694 | InactiveE_557 | GGTATGAATATCTAATAGTTGTGAATTGAAAAGTCTTGAC | ATTO647N |
| chr11 | 55833696 | 55833736 | InactiveE_558 | CCTACCTTAAGCCTTACCTTATGGTATGATCCTCACATCT | ATTO647N |
| chr11 | 55833738 | 55833778 | InactiveE_559 | ATTACGTTATAGCCTCCTGATTTTCACTCTGTACTATTAG | ATTO647N |
| chr11 | 55833780 | 55833820 | InactiveE_560 | AGACCATATGTTAGTTGCCACAGCCAGAAATAAATCACAG | ATTO647N |
| chr11 | 55833822 | 55833862 | InactiveE_561 | GCTCTCTGATACTCTTCATGTTAGGAGTAAGGCTATGGTT | ATTO647N |
| chr11 | 55833948 | 55833988 | InactiveE_564 | AAATTATTCATCTTTCACCTCTGTATCCACCTCAGATATT | ATTO647N |
| chr11 | 55833990 | 55834030 | InactiveE_565 | ATGGCTTTCTCTCTAACTAGAGTGTGGATTCCTGGAGATA | ATTO647N |
| chr11 | 55834032 | 55834072 | InactiveE_566 | CTCAGGTCACAGTGTGACGTAGCTATCTGCTTATACATTT | ATTO647N |
| chr11 | 55834074 | 55834114 | InactiveE_567 | GGGTACTCACAATTTCTGTGAACAGCCTGTATTTTATTGT | ATTO647N |
| chr11 | 55834116 | 55834156 | InactiveE_568 | TGAGACATTTTCTGACTCCTCCCAACCAAGTTCAGTATTT | ATTO647N |
| chr11 | 55834158 | 55834198 | InactiveE_569 | TTATCCATACAGCAAAACATTGCTGAGTTGTGAAAACTTC | ATTO647N |
| chr11 | 55834200 | 55834240 | InactiveE_570 | TGGCTCCCAACAAACTAGTCCACTGCTAAATTGTCATTTA | ATTO647N |
| chr11 | 55834830 | 55834870 | InactiveE_585 | ACAATAGGATTATTCCCCAGCATGTAAATCTTAGTGATTT | ATTO647N |
| chr11 | 55834914 | 55834954 | InactiveE_587 | TGTAAAACATGCAAATCTCACCAAACTTCATGAGTTGTGC | ATTO647N |
| chr11 | 55834956 | 55834996 | InactiveE_588 | AACTGCAACAATTTGGATAATTAACGTGTAATCAAATTGT | ATTO647N |
| chr11 | 55835124 | 55835164 | InactiveE_592 | GAGAAAGTGGACCATGTGAATATATCACTTATTCCAAAAT | ATTO647N |
| chr11 | 55835166 | 55835206 | InactiveE_593 | ATGCCTATACTGGGGAGTTTTTGCAGGGCTGATATAATAA | ATTO647N |
| chr11 | 55835250 | 55835290 | InactiveE_595 | TCACACTACCACTAAAATAACTCTTTACAAATAAATCTAC | ATTO647N |
| chr11 | 55835292 | 55835332 | InactiveE_596 | TTGGTCCTTATGCCTATAAATTTATCCTCTTATATATGCC | ATTO647N |
| chr11 | 55835334 | 55835374 | InactiveE_597 | CATTTTATAATTTAACACTTGCACTGTTCAATAGCCTCCA | ATTO647N |
| chr11 | 55835460 | 55835500 | InactiveE_600 | TCCTCTTTGGTTCTGATTTACCATATAAATTTCATCAAGC | ATTO647N |
| chr11 | 55835544 | 55835584 | InactiveE_602 | CTGATGTTGTTGCAAACATCCCACATATCCTGTAAACAAA | ATTO647N |
| chr11 | 55835586 | 55835626 | InactiveE_603 | CATAAGAAATTACGTCTAGCTACAACCAATATATTAAGGT | ATTO647N |

| | | | | | |
|---|---|---|---|---|---|
| chr11 | 55835628 | 55835668 | InactiveE_604 | GGACAGGACTGCAAACGTTTGGAATTTGTTTCTCAGTGTC | ATTO647N |
| chr11 | 55835670 | 55835710 | InactiveE_605 | GCCAAGGATATTAGATGCAGATCCGATCTTCATGTATAGA | ATTO647N |
| chr11 | 55835712 | 55835752 | InactiveE_606 | GAGATTCTTCCTTCAGGTCTCTCATCTGCTTAGTTAATTA | ATTO647N |
| chr11 | 55835754 | 55835794 | InactiveE_607 | TAACTGTTCTCTCTGTTACACTAACCACACAAGAGATGCA | ATTO647N |
| chr11 | 55835796 | 55835836 | InactiveE_608 | TATTTCTTATATATCTACCATGCCCTTGGTCTTTGCAAAT | ATTO647N |
